# Continental-scale hyperspectral tree species classification in the United States National Ecological Observatory Network

**DOI:** 10.1101/2021.12.22.473714

**Authors:** Sergio Marconi, Ben. G. Weinstein, Sheng Zou, Stephanie A. Bohlman, Alina Zare, Aditya Singh, Dylan Stewart, Ira Harmon, Ashley Steinkraus, Ethan P. White

## Abstract

Advances in remote sensing imagery and machine learning applications unlock the potential for developing algorithms for species classification at the level of individual tree crowns at unprecedented scales. However, most approaches to date focus on site-specific applications and a small number of taxonomic groups. Little is known about how well these approaches generalize across broader geographic areas and ecosystems. Leveraging field surveys and hyperspectral remote sensing data from the National Ecological Observatory Network (NEON), we developed a continental-extent model for tree species classification that can be applied to the network, including a wide range of US terrestrial ecosystems. We compared the performance of a model trained with data from 27 NEON sites to models trained with data from each individual site, evaluating advantages and challenges posed by training species classifiers at the US scale. We evaluated the effect of geographic location, topography, and ecological conditions on the accuracy and precision of species predictions (72 out of 77 species). On average, the general model resulted in good overall classification accuracy (micro-F1 score), with better accuracy than site-specific classifiers (average individual tree level accuracy of 0.77 for the general model and 0.70 for site-specific models). Aggregating species to the genus-level increased accuracy to 0.83. Regions with more species exhibited lower classification accuracy. Predicted species were more likely to be confused with congeneric and co-occurring species and confusion was highest for trees with structural damage and in complex closed-canopy forests. The model produced accurate estimates of uncertainty, correctly identifying trees where confusion was likely. Using only data from NEON, this single integrated classifier can make predictions for 20% of all tree species found in forest ecosystems across the entire US, which make up to roughly 90% of the upper canopy of the studied ecosystems. This suggests the potential for integrating information from multiple datasets and locations to develop broad scale general models for species classification from hyperspectral imaging.

## 1. Introduction

Forest ecosystems play a central role in essential services like providing wood and other forest products, carbon sequestration, and biodiversity conservation (Wiens, 2016; Pecl et al., 2017), but understanding patterns and processes driving forest properties and species distributions across scales can be challenging. A common strategy to monitor biodiversity and biomass of forests at national scales is to use field surveys of plots (USDA Forest Service, 2001, Lawrence et al. 2010). Data collection within survey plots requires extensive effort, limiting even the most extensive national forest inventories to several thousand permanent plots sampled every few years (White et al., 2016), which can be too sparse for investigating the effects of management, soil properties, topography and local environmental conditions on large scale forest structure, distribution and diversity (Tomppo et al., 2008). Remote sensing can help bridge this gap between local and regional scales by providing individual tree level data at scales beyond what is feasible for traditional plot-level inventories (Anderson, 2018). Models linking remotely sensed imagery to field surveys can identify the location and species identity of individual trees (Henrys & Jarvis, 2019), alleviating the challenge of inferring local patterns from sparsely sampled data (Ayrey et al., 2019, Bastin et al., 2019, Kandare et al., 2017) for understanding tree species distributions and abundances.

Numerous approaches have been developed for pixel- or canopy-scale species-level classification using hyperspectral remote sensing based on exploiting spectral differences between tree species which are caused by differences in foliar properties and canopy structure (Shi et al., 2018, Mayra et al, 2021, Belgiu & Dragut, 2016, Ballanti et al., 2016, Ab Majid et al., 2016). Recent efforts in species classification use either deep learning methods (Nezami et al,, 2020, Zhang et al., 2020, Martins et al., 2021) or ensemble of machine learning (Knauer et al., 2021, Grabska et al., 2020), showing promising improvements over more traditional approaches such as random forest, support vector machines or multi-layer perceptron classifiers. In general, most approaches are conducted with datasets covering small site-and/or ecosystem-specific extents (Fassnacht et al., 2016) rarely focus on classification of individual trees (but see urban tree mapping e.g. Martins et al., 2021), and often focus only on less than 10 species (Michałowska et al. 2021). For example, because of limitations related to coarse pixel size, many studies using satellite data either predict the dominant species within plot-sized pixels (Grabska et al., 2020, Wang et al., 2022) or classify the relative distribution of broad vegetation types within pixels (Bogan et al., 2019). These approaches are valuable for addressing processes for which information about dominant species in the community or ecosystem type is needed (e.g. monitoring forest aboveground biomass, Laurin et al. 2020), but are currently limited in their ability to provide precise taxonomic information at the individual level. Precise fine-grained species information is important for assessing forest biodiversity, tree-level growth and species interactions (Anderson, 2018). Other recent works have leveraged high resolution airborne missions to generate tree surveys covering hundreds of km^2^ and encompassing multiple management regimes and forest types (Modzelewska et al., 2020, Modzelewska et al., 2021). Yet these works target single biomes, and so even though they provide valuable surveys for key species across different stand ages, communities structures and topographic positions, their use is still limited to individual biomes and relatively small regions.

Developing remote sensing models specifically for individual regions, sites and/or ecosystems, as is typically done with remote sensing from airplanes and UAVs, limits the use of the models beyond the region and training data, making it difficult to: 1) conduct research at regional to continental scales due to the lack of general models that can be applied across ecosystems; 2) identify rare or uncommon species due to limited data for training models, which often results in studies focusing on a limited subset of common species; and 3) accurately apply the model beyond the region or conditions of the associated field data. Furthermore, training data from single site studies often lack the full range of variation in spectral characteristics that can occur for each species due to intraspecific variation. Developing generalizable species classification models based on data across different forest types and large spatial extents unlocks the potential for overcoming these limitations and increases the utility of remote sensing for building reliable broad scale tree species surveys.

Developing individual tree level species classification models that span geographic areas, forest types and species pools poses a novel set of challenges. First, it requires building a library of co-registered field and remote sensing data that includes data from multiple sites and ecoregions for training and testing algorithms. Second, increasing the geographic extent of species classification risks confusing species that have similar spectral properties but do not overlap in their geographic distributions. Third, combining data from multiple sites may introduce variation in spectral reflectance due to differences in phenology (which affect leaf greenness) and environmentally driven intraspecific variation, which affect leaf biochemistry, crown shape and leaf biophysical traits (Sims & Gamon, 2002). Finally, aggregating remote sensing data from multiple flights, sensors, and sites may increase variation in spectral signatures due to complex sources of spatial and temporal variation that are linked, but not limited to, acquisition dates, solar angles, ecosystem types and variation in sensor calibrations (Pax-Lenney et al., 2001). Therefore, while there are many potential benefits to models for species classification across large spatial extents, it is unclear how they will perform compared to local models developed for specific ecosystems.

Here, we leverage newly available data from the National Ecological Observatory Network (NEON) to develop a continental level model for tree species classification that can be applied to the entire network and compare its performance to the traditional approach of building individual models for each site. We used NEON remote sensing and field data on individual trees at 27 terrestrial sites from Puerto Rico to Alaska, covering a wide range of ecoregions and biomes across the United States (US). Several studies have developed species classification models for NEON data, but all these studies focused on individual NEON sites (Scholl et al., 2020, Fricker et al., 2019, Marrs & Ni-Meister, 2019, Marconi et al., 2020), or 2-3 sites in the same region (Graves et al. 2021). We build on these single site models to develop a general model that can be applied across the entire NEON network by connecting field-identified tree stems to hyperspectral images. We used an ensemble of species classification models to allow for leveraging the strengths of different machine learning classifiers and provide effective ways to estimate the uncertainty of predictions (Engler et al., 2013, Saini & Ghosh, 2017, Sagi & Rokach, 2018). Using this model, we (1) assess whether a general model approach improves performance compared to separate models for each site, (2) determine the importance of reflectance, geography, environmental and ecological conditions on the accuracy and precision of species predictions; (3) evaluate the uncertainty in predictions; and (4) discuss the potential for this general model to be used for ecological applications.

## 2. Methods

### 2.1. Field Data

Vegetation structure field data (https://data.neonscience.org/data-products/DP1.10098.001) were collected by the NEON terrestrial observatory system (TOS) between 2015 and 2019 (Table S.1). This dataset, sampled from 400 m^2^ plots distributed across the landscape of each NEON site, includes information about individual trees’ geolocation and properties such as species identity, health status, canopy position, crown diameter, and tree height. Vegetation structure plot locations are located randomly across the sites stratified by vegetation type within each site with the aim of capturing landscape level biological and structural diversity at each site. Each subplot (200m^2^ in size) is assigned to an ecosystem type extracted from the National Land Cover Dataset. For this study we used data from 27 of the 41 NEON sites with partial to complete forest cover, encompassing 17 out of 18 ecoclimatic domains in the US (Figure S.1). We used a total of 1701 subplots from 714 plots. Data from the other NEON sites could not be used because either field data about tree stem positions was missing or the remote sensing imagery contained gaps in the hyperspectral or lacked information about the sensor angle at the time of data collection. We only included individual stem data that met the following criteria: (a) the stem had a species label assigned to it, (b) it was marked as “alive” and “tree” in the NEON field inventory, and (c) it belonged to a species with more than 5 entries for the entire cross-site dataset. We also did not use stems designated in the NEON vegetation structure data as fully shaded, shrubs or sapling, as these stems are most likely not visible in the remote sensing imagery and would therefore be erroneously paired with pixels belonging to species from neighboring overstory crowns. The final dataset used for species classification consisted of 5697 individual trees of 77 species.

### 2.2. Remote sensing data

For this study we used the hyperspectral L3 data from the NEON Airborne Observatory Platform (NEON, 2021). These data are provided in 1 km^2^ tiles with 426 channels recording reflectance in 5 nm bands from 350 to 2450 nm. Reflectance data was atmospherically corrected using the ATCOR-4 approach (Krause et al., 2011). Pixel size is 1 m^2^. We applied bidirectional reflectance distribution function (BRDF) correction, topographic correction, and L2 normalization to reduce the effect of peripheral light and non-Lambertian scattering with the goal of minimizing variation in reflectance ascribable to flight path and airplane position (Marconi et al., 2020). For all tiles (n = 4500), we used the same general parameterization to define the BRDF kernel. We also dropped bands in the water absorption regions of the spectra (1340 – 1430 nm and 1800 - 1955 nm) as well as the spectrometer’s peripheral bands to reduce the effects of noise and artifacts. Thus, the hyperspectral data were reduced to a total of 347 channels. In the tree species classification models, we included terrain elevation (1 m^2^ spatial resolution) along with the hyperspectral data because of elevation’s potential information in discriminating species within landscapes (Strahler et al., 1978, Scholl et al., 2020). Elevation data were derived from a LiDAR sensor mounted along with the hyperspectral sensor on the aircraft, which was converted into a 1 m spatial resolution raster and appended to the hyperspectral data as an additional band.

We assigned each individual tree from the filtered field dataset to a square clip of 16 pixels (4 m crown diameter), centered around the stem’s GPS coordinates. This threshold was selected because it is smaller than more than 95% of individual tree crowns diameter measured from the NEON vegetation structure dataset. We adopted this strategy to reduce the number of mislabeled pixels at the edges of the crown that belong to neighboring trees, especially in dense closed canopies. To remove shaded and non-vegetation pixels from these clips, we removed all pixels with NDVI < 0.5 and low reflectance in the NIR (reflectance at 825nm < 0.2). Since stem positions often do not match precisely with the center of the tree crown in the canopy, pixels will sometimes be assigned to the wrong label. To reduce this, we filtered out pixels that were much shorter than the maximum height of the crown. These pixels are less likely to belong to the sunlit portion of the target crown or may even measure the reflectance from neighboring shorter tree crowns, or the understory within a gap in the target crown. We filtered out pixels that were ≥5 m below the top height of the tree as determined by the maximum height of the tree from the LiDAR data in the 16-pixel clip. Finally, we removed stems with field GPS locations that fell within 3 meters of one another where the stems belonged to different taxa to decrease the chance of confusing closely neighboring, and potentially intermixed, tree crowns of different species. After all these steps, the final, filtered dataset used ∼50,000 out of 200,000 initial pixels and 6449 out of ∼21,000 crowns in the original vegetation structure dataset.

Due to the large number of correlated bands in hyperspectral data, it is necessary to reduce the number of features used in classifiers and limit the potential for overfitting (Li et al., 2011). Although PCA is the most common approach to achieve dimensionality reduction, it comes with a number of limitations that could be problematic when aggregating information from different image collections, since it is sensitive to outliers, assumes linear relationship across features, and it is prone to discarding low rank components that may have high discriminative information (Prasad & Bruce, 2008). An alternative solution to reduce these issues is to use untransformed hyperspectral reflectance and group highly correlated bands based on their distribution in the form of probability densities (Delicado, 2011). This is possible using a hierarchical dimensionality reduction, consisting of clustering bands with similar standardized distributions according to Kullback-Leibler divergence (KLD) (Zare et al., 2019). The advantage of this approach is that it allows for reducing the number of features used while using untransformed spectral information, thus identifying redundant bands, highlighting highly correlated regions of the spectra (Yang et al., 2014), and allowing for a direct identification of the most informative spectral regions. The main limitation is that it requires arbitrarily choosing the number of groups into which to cluster the bands and identifying meaningful summary statistics to summarize the information clustered in the groups. We chose 15 groups of bands because given the limited number of individuals available per rare species, a smaller number of features is necessary to minimize model overfitting on train data. The number of groups was selected after exploring a range of possible values from 8 to 40. Fewer groups resulted in a loss of information and generally lower accuracy, while more groups did not significantly change model performance. Groups of bands were trained using pixels in the training data. Since the KLD clustering resulted in grouping bands from mostly contiguous and distinct spectral regions (though on the boundary of some groups of bands the bands put into each group was discontinuous), we chose the maximum, minimum and average reflectance as features to measure the peak of reflectance, peak of absorption and average reflectance within each spectral region, which have been linked to leaf traits and vegetation properties (Artiola et al., 2004). This allowed us to reduce the 347 hyperspectral bands into 45 distinct features quantifying including information on the minimum, maximum and mean for each of 15 spectral regions (i.e., groups of bands).

### 2.3. Site effects

To provide the model with information on site location, which could reduce confusion across species that do not co-occur within a site but are characterized by similar spectral signatures, we included the latitude and longitude of the centroid of each site in the model. This approach incorporates information on the proximity of different sites and can be readily generalized to use outside of NEON. To help control for potential differences resulting from variation in sensor calibration of the specific flight missions, which would be specific to each site, we added a “site identifier” to the remote sensing features in the model. The site identifier consisted of the NEON site names (a nominal variable) transformed into real positive numbers by applying Leave-One-Out regression encoding, based on the correlation between the categorical variable (i.e. site name) and the species classes for each site(https://contrib.scikit-learn.org/category_encoders; Wright & König, 2019). The advantage of this approach over the more commonly used one-hot-encoder (i.e., adding a binary feature for each site in the dataset) is that it compresses the information into a single feature, which avoids undesired sparsity and potential overfitting due to a large number of encoded classes (27 in this study) (Rodriguez et al., 2018). We used data in the training set to fit the encoder and assigned its average value to each site category. The final model input for the general model was hyperspectral features, elevation, latitude and longitude and site. For the site-specific models, only spectral features and elevation were used.

### 2.4. Species classification

To assess whether a general model approach improves performance we built two sets of models: (1) a general model using data from all 27 NEON sites and (2) 27 separate models, each one using only the data from a single NEON site and covering a region of few hundred km^2^ (hereafter referred to as site-specific models). For both the general and site-specific models, we performed species classification at the pixel level using an ensemble of five classifiers (Figure S.2): (1) a random forest classifier (Belgiu & Dragut, 2016), (2) a k-nearest neighbors classifier (Laaksonen & Oja, 1996), (3) a histogram gradient boosting classifier (Guryanov, 2019), (4) a fully connected multilayer perceptron (Pacifico et al., 2018), and (5) a bagging classifier with support vector machine as base estimators, using tools from the scikit-learn python package (Pedrosa et al., 2011). Details for each classifier can be found in supplementary materials (Supplement 1: classifiers). Ensemble-based approaches generally provide better performance and limit overfitting compared to using one classifier alone (Knauer et al., 2019). We chose the individual classifiers which form the ensemble because they have been shown to perform well for species classification on NEON data (Marconi et al., 2019). All predictors were normalized for model fitting by subtracting the mean and dividing by the standard deviation (i.e., setting the mean to zero and the standard deviation to 1). Parameters for all models and the ensemble were extracted by performing parameter tuning using cross validation.

We used entropy loss to measure the quality of tree-splits for random forests, categorical cross-entropy as the loss function for the histogram-gradient boosting, a radial basis function kernel to allow for a non-linear decision surface for the support vector classifiers, and the Manhattan distance for calculating the distance between k-nearest neighbors in the KNN classifier. We stacked these five pixel-based models by using the probability vectors produced by each classifier as features for a meta-ensemble elastic-net logistic model (Tang et al., 2015, Hui & Hastie, 2005). We chose this approach because logistic classifiers are easily interpretable and use maximum likelihood to obtain estimates of the coefficients, returning as a result confidence scores that match the probability of a label-match and not just the single best predicted classification (Maddala, 1986), which is fundamental for assigning a reliable uncertainty score to each prediction. Pairing predictions to robust estimates of uncertainty is fundamental to increase the utility of remote sensing tree surveys for ecological analysis because it allows for (1) selecting trees and areas that meet or exceed minimum confidence in the derived measures for being used for scientific analyses, and (2) allows for cascading the uncertainty in predictions onto the results of analyses downstream (Dietze, 2017). Training the logistic meta-ensemble on calibrated scores from sub-classifiers offers an advantage over other modern algorithms, whose estimates of uncertainty do not match true probabilities and are not well calibrated to the output of interest (Guo et al., 2017, Mukhati et al. 2020).

One of the main challenges of species classification algorithms is the imbalance between number of individual samples for rare and common species, which can cause models to overfit to highly abundant classes. In our data set, the number of pixels per species ranged from 44-28000 and the number of individual trees per species ranged from 5-1000. We used SMOTETomek technique (Batista et al., 2003) to reduce the effects of species class imbalances in the training set. SMOTETomek consists of a combination of under and oversampling which resulted in roughly 1000 spectral signatures (pixels) per species. First, we undersampled pixels from the most abundant species using Tomek links, which removes noisy and borderline pixels (Tomek, 1976). Then, we used a SMOTE oversampling approach (Chawla et al., 2002) to create non-identical synthetic pixels for any species with fewer pixels than the majority class, thus balancing each class to roughly 1000 pixels each. No over-undersampling was applied to the test data. Because of the stratified design of the train-test split, most species and sites had a number of test trees proportional to their frequency in the original dataset. We also used the same train-test split to repeat the entire analysis once for each NEON site by building and testing site-specific models built using only data from each particular site. Finally, to estimate which spectral regions are most important for separating conifers from broadleaf species, we repeated the entire analysis by substituting species with broader taxonomy classes (i.e., angiosperms vs gymnosperms).

### 2.5. Evaluation

We evaluated the performance of the models by training the model on 80% of the data and evaluating its performance on the remaining 20%. Since spatial autocorrelation across train and test data can lead to optimistic bias in classification (Millard & Richardson 2015), we placed all individuals within a plot together into either the training or testing data sets. A series of randomizations of the plots were performed to create an 80:20 split of individuals that optimized the number of species in the train and test data sets. For each randomization, we calculated the total number of species in the test set and repeated this random operation until we found the split which maintained the highest number of species from the original data in both train and test set. For the general model, the training data set contained 4210 individuals of 77 species and the test data set 1487 individuals of 72 species. Data for the 5 species missing from the test-set were collected only within plots selected for training, therefore no tree from these 5 species was suited or included in the held out testing data to minimize the effect of geographic autocorrelation on assessing accuracy (Karasiak, 2021). The resulting data represents 56% of the total tree species in the original unfiltered vegetation structure dataset and these species account for an average of 89% of individuals per site (Figure S.3).

Predictions for the species class of each individual in the test set were made using a 4×4 clip centered on the location of the test stem. Model performance was then evaluated using overall accuracy, individual tree level (micro) and average species-level (macro) F1 scores (hereafter referred to as individual-level and species-level accuracy respectively). The F1 score species classification, allowing for direct comparison between models using a single metric (Chinchor, 1992). For each site, F1 scores for the general model were compared to those produced by equivalent single site models to determine how the general model performed relative to the traditional single site approach. Scores and confusion matrices were calculated using the Caret package (Kuhn, 2008).

To understand the performance of the general model in different ecological contexts, we evaluated how performance varied across the United States, how performance correlated with the number of species being predicted at the NEON site, and which components of the model (site effect, elevation, geographic location, and hyperspectral reflectance) were most important for prediction. We used bootstrap features importance to quantify the relative importance of the different types of features, e.g., site identifier, site geolocation, hyperspectral reflectance and terrain elevation (Breiman, 2001). This approach is based on evaluating how the overall accuracy is affected by each individual feature. At every iteration, one feature is selected and the values are randomly shuffled among the samples, effectively removing the information held in it. The accuracy is recorded with the shuffled feature to determine the loss of performance compared to the unshuffled data. We used the same approach to quantify the relative importance of the 15 spectral regions in which we grouped the hyperspectral data.

We also evaluated the characteristics of trees and forests associated with the most confusion between species (i.e., misclassification) based on forest type (using the National Land Cover Database; Homer et al., 2001) and information from the NEON field data on canopy position, tree status, and growth form from the NEON field data. We also assessed spatial structure in confusion by determining, for every misclassified tree, whether the species to which it was incorrectly classified to also occurred in the same NEON field plot. Finally, since confusion commonly occurred within genera we also evaluated model performance for predicting genus instead of species.

### 2.6. Prediction

We generated predictions for individual trees at the landscape scale (∼350 km^2^) by integrating our approach with individual tree detections from previous work (Weinstein et al., 2021). The Weinstein et al. (2021) dataset consists of 100 million individual tree crowns from 37 NEON sites identified using a retinanet neural network object detector and represented by quadrangular polygons (i.e., bounding boxes) roughly representing the surface of the sunlit portion of the crown. For consistency between the approach used for training and testing the model (16-pixel clips), we extracted the pixels from the centroid of each estimated bounding box. First, we extracted a 4×4 square window of pixels around the centroid of each detection. For bounding boxes smaller than 16m^2^, we dropped the pixels falling outside the bounding boxes. Second, we filtered vegetation pixels from the background using the same procedure as applied to the training/test data set. We finally selected all pixels with uncertainty scores > 0.5 to be used to make predictions at the level of individual trees. We assigned each tree to a species class by averaging the probability vectors (i.e., probability that the pixel is assigned to any of the 77 classes) of each pixel in the crown and selecting the species with the highest average probability. We assigned each individual-tree prediction an uncertainty score consisting of the average pixel probability, which ranged from 0-1.

## 3. Results

The general (cross-site) model yielded more accurate species classifications (larger F1 scores) than site-level models for 13 (species-level F1) and 18 (individual-level F1) of the 27 sites and identical accuracies for 5 (species-level F1) and 6 (individual-level F1) additional sites. There were only three sites that showed better site-level species-level and individual-level F1 scores: Blandy Experimental Farm, Washington (BLAN) and Talladega National Forest, labama (TALL), and Jones Ecological Research Center, Georgia (JERC) (Figure 1, Figure S. 4, Figure S. 5). On average, the general model resulted in higher accuracy of individual tree level classification (increases in individual-level F1) from 0.70 to 0.77 and species-average accuracy (increases in species-level F1) from 0.46 to 0.54. Accuracy of the ensemble was higher than its sub-models trained singularly whose average site-level accuracy ranged between 0.09 and 1 species-level F1 and between 0.31 and 1 individual-level F1 (Figure S.7 and Supplement 2 for detailed species level accuracies, site-level and general model confusion matrixes in Supplement 3, raw outputs available at https://doi.org/10.5281/zenodo.5796142), which is consistent with the general observation that ensemble-based approaches produce more accurate predictions (Healey et al., 2018). Since the general ensemble model proved to be the best performing approach in this study, we focus primarily on it from this point forward.

**Figure 1.**
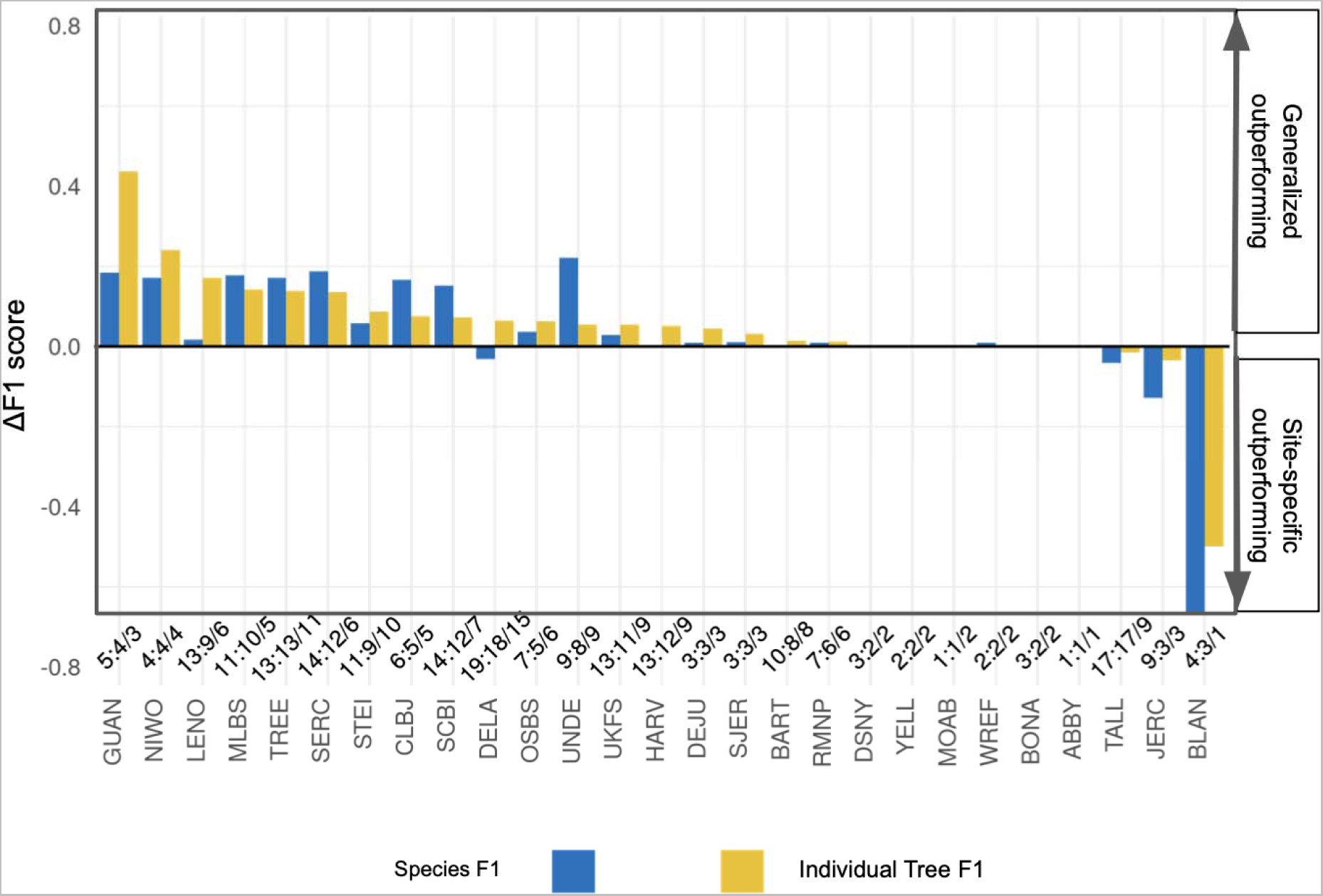
Performance of generalized vs site-specific classification models for each NEON site. Positive values are sites for which the generalized model performed better than site-level. Negative values are sites for which the generalized model performed worse compared to site-level. Blue bars represent species-level F1 score, yellow bars individual-level F1. Numbers separated by (:) on top of each site name represent the total number of species in the training for each site (general model: site-only model).

Our results show a link between classification accuracy and ecological properties such as ecosystem type, tree health, and growth form (Figure 2, Figure S.6). Damaged trees, including broken boles and other types of damage (but not diseased trees), exhibited higher rates of misclassification than healthy crowns (Figure 2), with broken boles exhibiting a 44% misclassification rate. The general model performed best in evergreen forests (∼12% misclassification rate) and worst in wetlands (∼38% misclassification rate), with deciduous forests falling in between (∼30% misclassification rate). Average classification accuracy was higher in eastern forests compared to western forests (Figure 3a), and was negatively correlated with the number of species within the site (Figure 3b,d). The algorithm generally underperformed in the Prairie Peninsula and Central and Southern Plains ecoregions which are characterized by patches of closed forest at the edges of prairies or farmland (Figure 3a, Figure S.6). These results align with previous work in showing that classification from remote sensing is more challenging for more complex canopies, overlapping crowns, and coexisting species with similar life history and spectral properties (Heinzel & Koch 2016, Bioucas-Dias, 2013).

**Figure 2.**
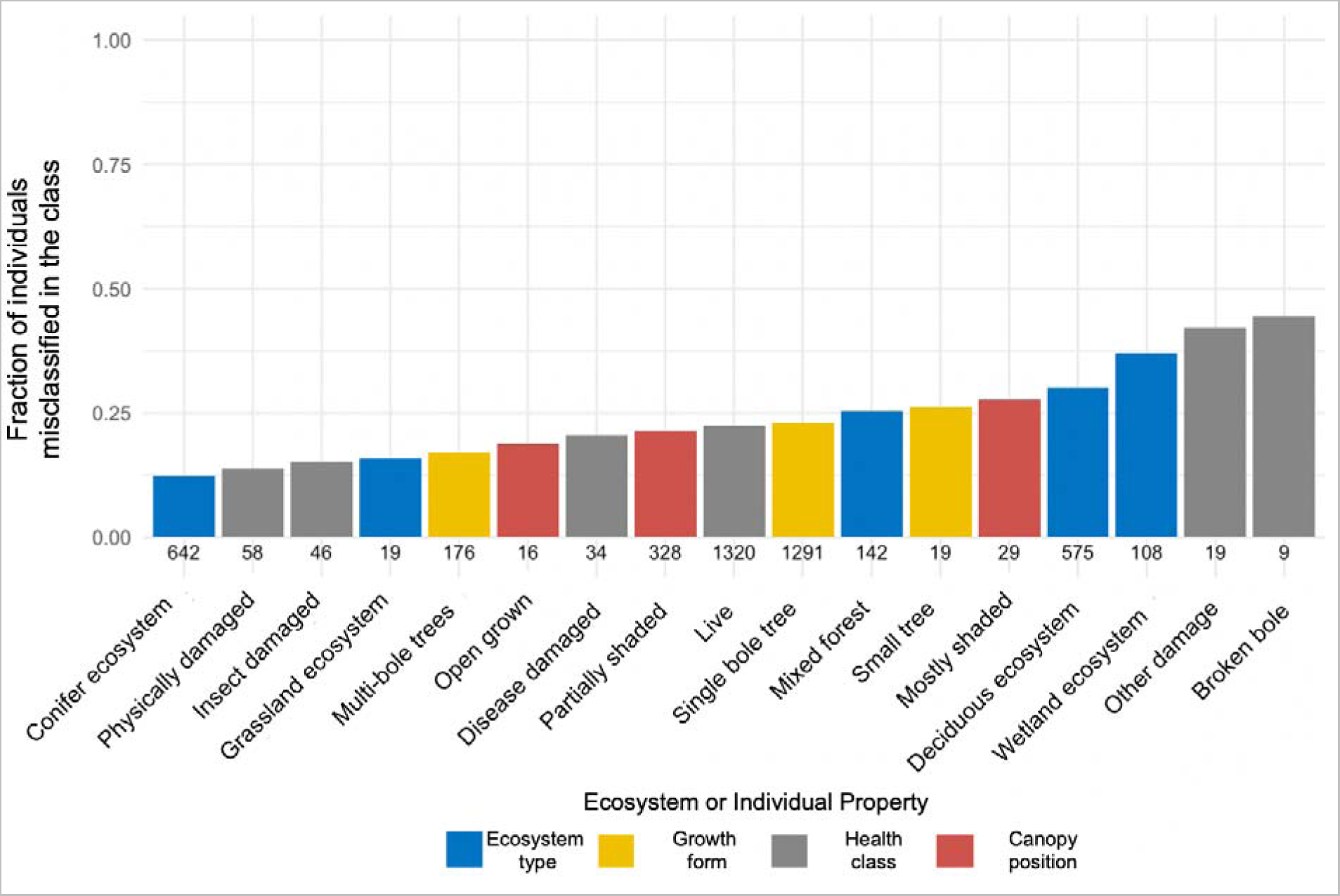
Fraction of misclassified trees across ecosystem types (blue), growth form (yellow), canopy position (red) and health status (gray). Numbers above the x-axis labels are the number of trees in each category.

**Figure 3.**
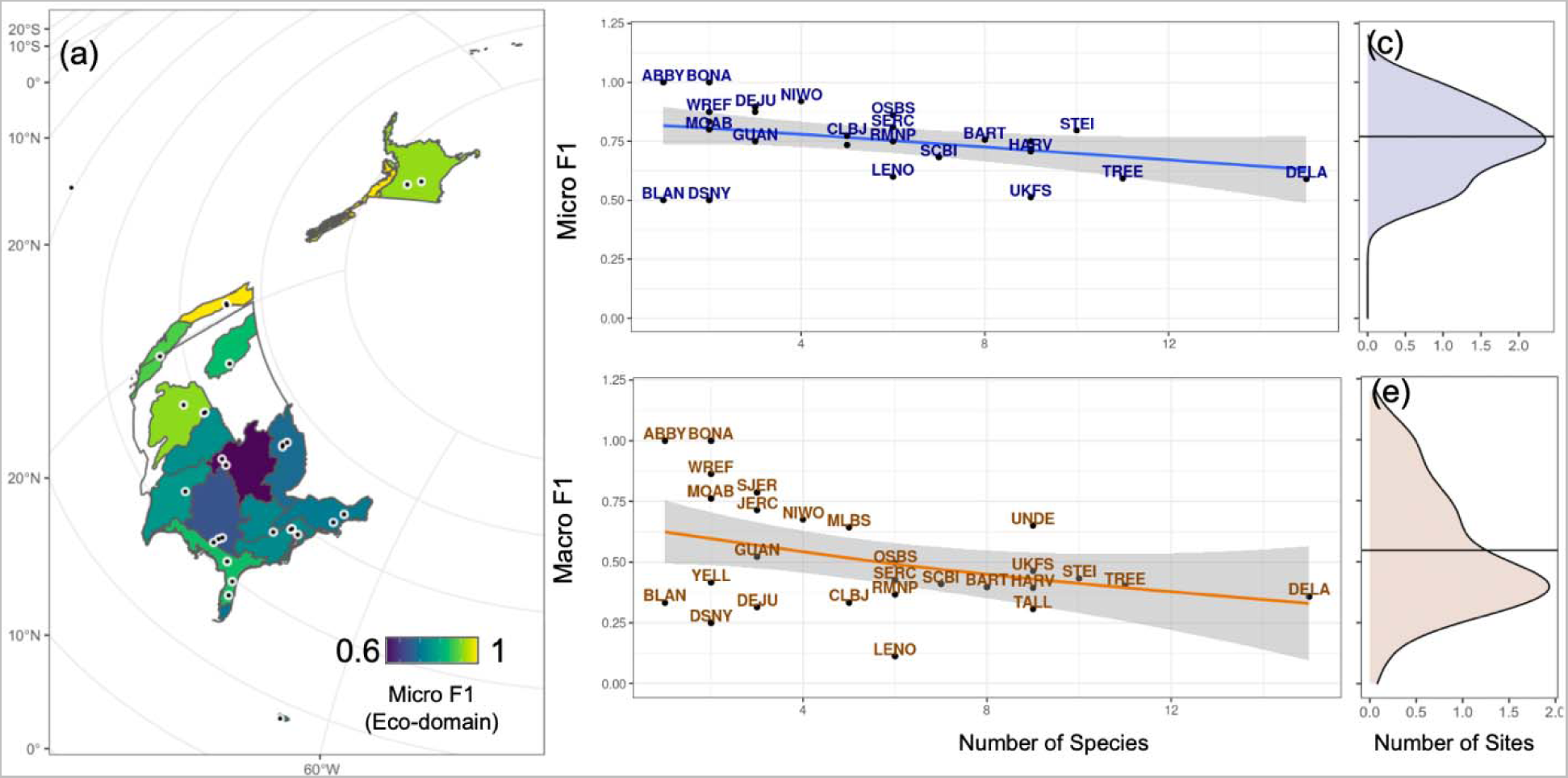
Variation in accuracy of the generalized algorithm across the US. (a) map of average individual-level accuracy (Micro F1) for each ecological domain. Dots represent the location of each NEON site. Blue polygons represent the Prairie Peninsula and Central-Southern Plains. (b) Relationship between individual-level accuracy (Micro F1) and number of species in the training dataset for each site (Number of Species). The blue line is the loess smoother relationship over the 27 sites. (c) Kernel density estimate of the distribution of individual-level F1 scores (averages per site). (d) Relationship between species-level accuracy (Macro F1) and number of trained species found in site (Number of Species); orange line is the loess smoother relationship over the 27 sites. (e) Kernel density estimate of the distribution of species-level accuracy scores (averages per site). Horizontal black lines in (d) and (e) represent the average accuracy across sites.

Roughly 80% of the information used by the algorithm for classifying species was from the hyperspectral reflectance (Figure 4). Important information was present across the entire spectrum, but our results show that some groups of bands in some spectral regions were more informative than others. Specifically, the most important spectral regions are the blue and green (0.450 to 0.550 nm) in the visible region, the red-edge in the near infrared (0.62 to 0.85), 1.15 to 1.27 nm in SWIR1 and 1.62 to 1.68 nm in SWIR2. Spectral regions in the SWIR1, SWIR2, and red-edge were the most important also in classifying angiosperms vs gymnosperms. The site’s coordinates, which represent the geographic locations of sites, explained 11% of total variation and were the second most important variable (Figure 4). Elevation, a proxy of potential local changes in the environment within each site, accounted for another 4%. The site effect, a proxy of other site level ancillary information (e.g., sensor calibration, flight and atmospheric conditions), only accounted for 3% of the total explained variance.

**Figure 4:**
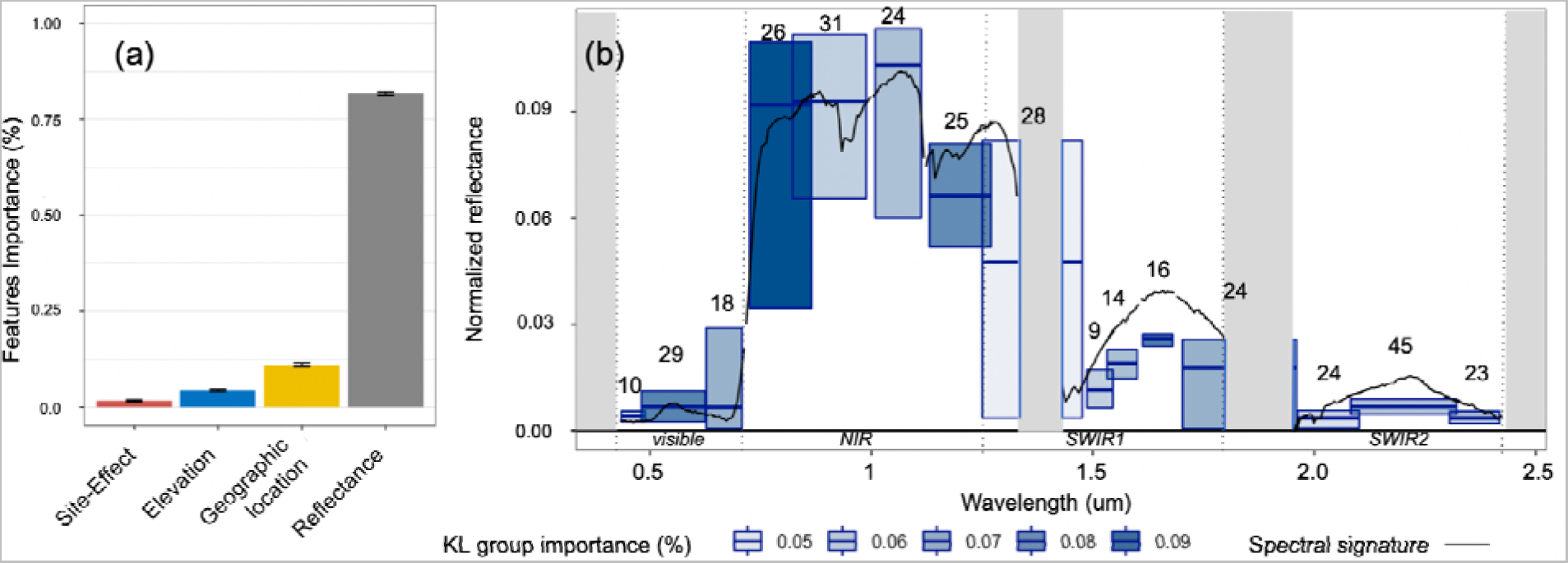
Features importance calculated from the permutation feature importance procedures described in Breiman, 2001, on the meta-ensemble model. (a) Relative contribution of different feature types: reflectance, as the sum of the 45 features (gray), site coordinates (yellow), elevation (blue) and site effect (red).(b) Relative importance of each Kullback-Leibler group of features used for dimensionality reduction of reflectance. Blue bars represent the reflectance for the average minimum, mean and maximum band in the specific KL group. Numbers on top of each bar represent the number of bands in each group. Bar width represents the range of bands covered by the specific KL group. Some bars overlap due to discontinuity of band assignments to different groups/bars at the group boundaries. Gray bars represent areas with water absorption bands dropped from the original hyperspectral images. Color intensity represents the relative importance of the specific KL group for the classifier (from light blue being of little importance, to dark blue being highly important). Black lines represent the reflectance of a randomly selected pixel to illustrate a typical vegetation reflectance pattern. Reflectance was normalized using L2 normalization. Numbers on top of each blue bar represent the total number of bands in the group.

Comparing misclassification among species shows there is greater confusion for rare species, congenerics, and species that co-occur within NEON field plots, and that model-estimated uncertainty accurately reflects confidence in the model prediction. All species performing poorly (F1 < 0.5) belonged to taxa with low sample sizes (less than 50 trees for training) (Figure S6, Figure S7). In general, most of the confusion was among species co-occurring within plot (74%) and site (93%). A large amount of confusion also occurred among congeneric species (∼27% of total misclassifications), mostly within pines, poplars, oaks and maples, which make up 57% of the test dataset (Figure S.9). Oaks, pines and poplars in particular accounted for ∼87% of the total within-genus confusion, and most misclassifications had confidence scores >0.8. Aggregating predictions at the genus level improved the overall accuracy by 6% (individual-level F1 accuracy of 83%), confirming that part of the confusion is embedded in physiological similarities across taxonomically related trees. Likewise, reducing tree classification into 2 plant functional types dramatically increased accuracy (F1 ∼0.95). The model showed a fair ability in predicting 5 of 9 species tested in sites where no data was used for that particular species in the training set. For these trees, the average individual-level F1 of ∼0.69 and average species accuracy of 0.47, but accuracy varied largely across taxa, with better results for needleleaf species (individual-level F1 ∼0.825, species-level F1 ∼ 0.71) compared to broadleaf species (individual-level F1 ∼0.44, species-level F1 ∼0.27). The model produced reliable estimates of uncertainty for all species regardless of the accuracy. Uncertainty scores matched closely with the probability of correct classification (R^2^ =of 0.89, Figure 5). Leveraging crown-data predictions, the model was tested to produce fair species predictions for millions of trees per NEON site (Figure 6).

**Figure 5.**
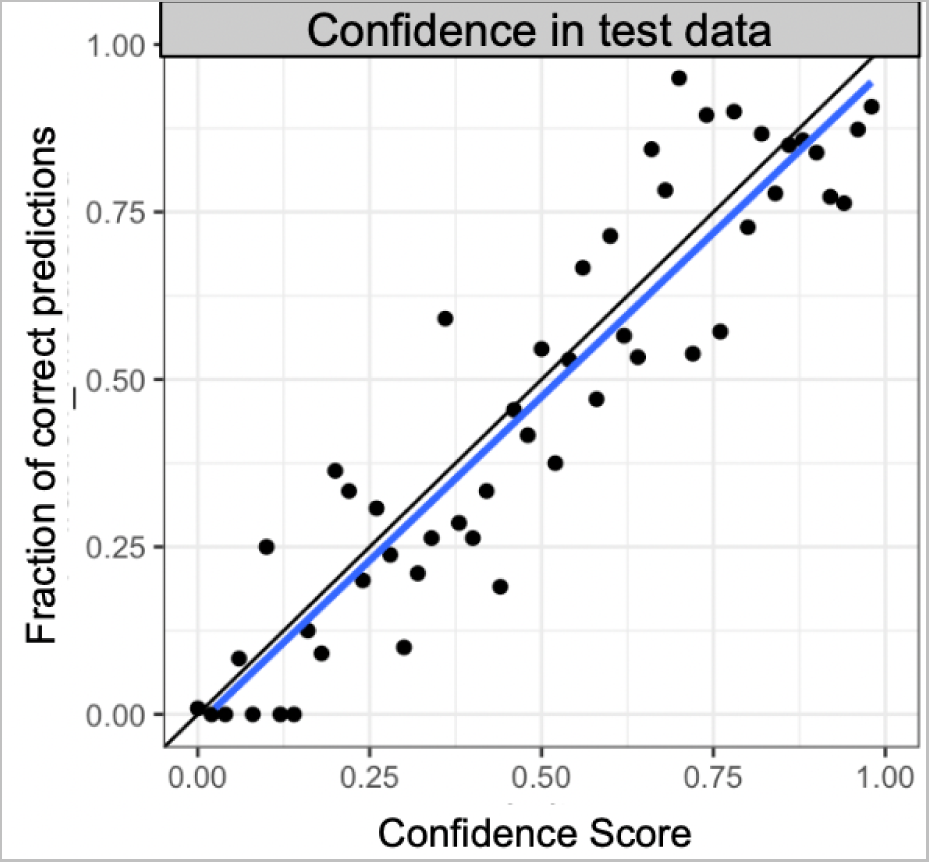
Evaluation of model confidence score (the probability of assigning the correct label to a prediction) as a measure of uncertainty. Confidence score was binned into 34 equal-width bins (each bin representing an interval of 0.03). Bin centers were plotted against the fraction of trees in that confidence score bin that were correctly classified. The blue line shows the fitted linear relationship between the confidence score and the proportion of correctly classified trees. The black line is the 1:1 line.

**Figure 6.**
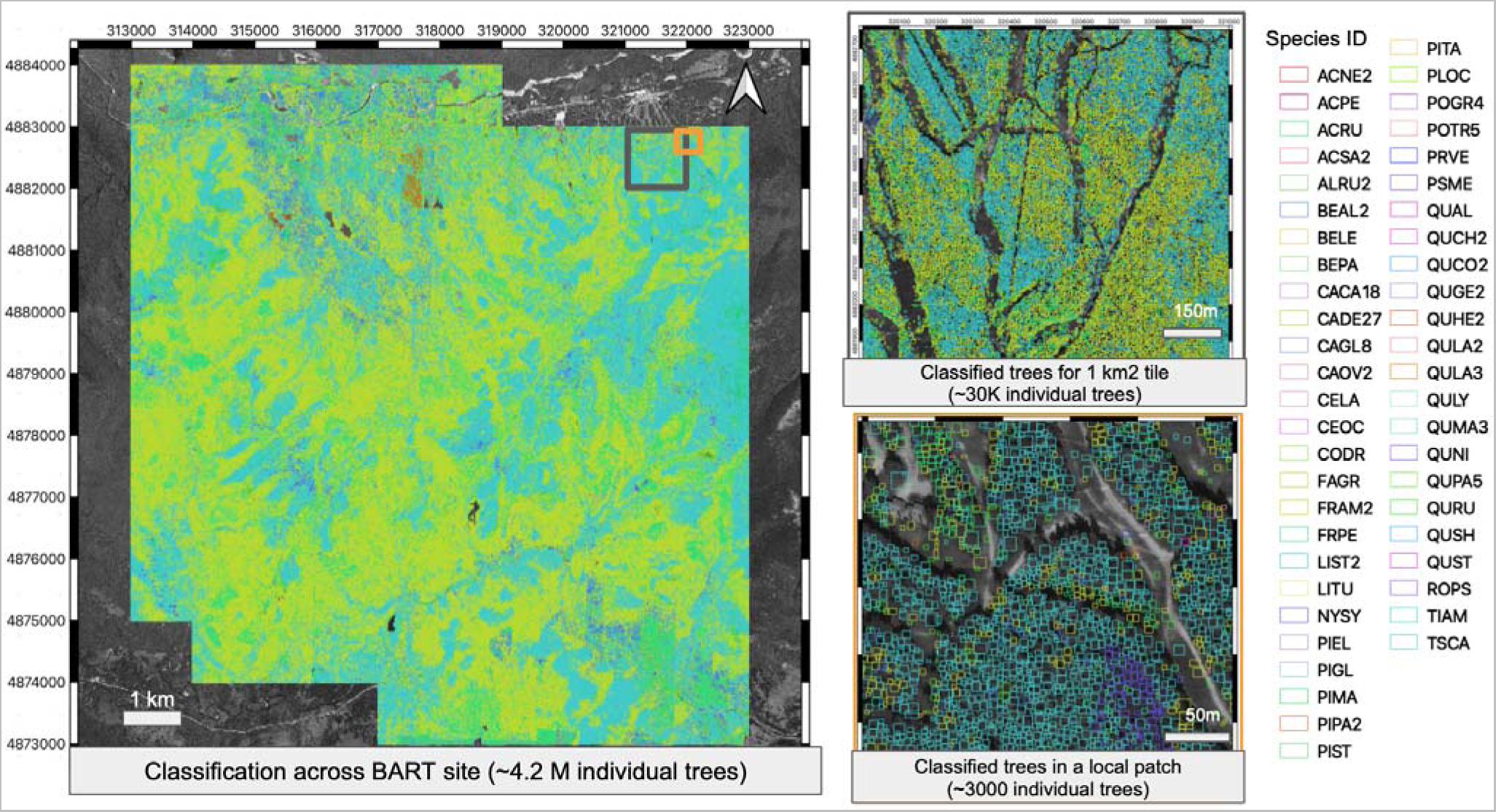
Example of species classification maps for all individual trees at the Bartlett Experimental Forest (BART) NEON site in New Hampshire. Species in legend include six of the most abundant taxa predicted at the site. Individual crown boundaries were estimated using predictions from Weinstein et al., 2021. The background is gray scale imagery of the site, so the gray areas on the left panel are regions for which NEON airborne data was not available; the gray areas on the right hand panels are areas without any trees including roads and other open areas.

## 4. Discussion

Using a single general model that integrated data from plots across a continental scale resulted in more accurate classification of tree species identity from remote sensing data than building separate models for individual sites. The more accurate classification occurred despite the continental data set containing samples from many different forest types, structures, and species compositions across 27 sites. This suggests that the benefits of increasing the number of samples for less common species and more fully characterizing within-species variance outweighs the costs associated with including species that do not overlap geographically and including components of within-species variance not observed at individual sites (Figure 1). To our knowledge this is the first study which developed a generalized model for species classification of individual tree crowns across multiple biomes. The success of the general model here suggests that developing generalized algorithms offers a potential step forward in species classification from remote sensing more broadly. Our model resulted in better cross-site classification compared to other approaches in literature (e.g., Castro-Esau et al. 2006) possibly because of the wider spectral range available from NEON hyperspectral images (445 - 2500 vs 445 - 950 nm), as suggested by the strong contribution of reflectance from 950 to 2500 nm to our generalized model (Figure 4). Also, better cross-site transferability of species classification may be related to the models used in the ensemble. Our model included methods like the gradient boosting classifier, which proved to be among the most robust for cross-site transferability of species classification (Graves et al., 2021). Our generalized approach leveraged the information from multiple locations, biomes, and survey efforts, increasing the number of individuals from rarely sampled or highly variable classes and allowing models to learn more broadly about how to distinguish species in the taxonomic group of interest. In addition to yielding improved predictions, generalized cross-site approaches can potentially generate predictions for a wide range of ecosystems, including those with limited or no training data, allowing other studies to leverage the same shared model and thereby facilitating large-scale ecological research (Weinstein et al. 2021).

By providing classification of the most common tree species in the canopy, the results of this model are potentially useful for several ecological applications, such as mapping biomass and modeling carbon, energy and water flux. Our model included species making up ∼80% of the individual trees in the upper canopy when all sites are taken together. The fraction, however, varied among sites. Furthermore, given the stratified sampling of the NEON vegetation structure data used to develop the generalized model, the model is likely to capture the major vegetation types and most common species at each site. Canopy trees, which are visible from optical remote sensing devices, represent the majority of biomass in forests (Lutz et al., 2012). Because they form the interface between the atmosphere and land surface, the canopy layer also is particularly important for water and energy flux (Paul-Limogens et al., 2017). Because carbon storage, water and energy flux can vary among species (Wright et al., 2006), the ability to map the location and coverage of canopy species is important for assessing these important ecosystem characteristics. Other ecological applications, such as assessing total forest species richness, and quantifying tree regeneration, cannot be addressed using the model because our model could not classify rare canopy species or understory individuals, One of the main challenges in developing models that generalize well across the continent is overcoming differences across sites in factors including seasonality, background, and sensor calibration (Hesketh & Sanchez-Azofeifa, 2012, Clark et al., 2005, Pu, 2021). To quantify the sensitivity of the algorithm to this ancillary information, we evaluated the relative importance of the site-effect features compared to reflectance, geography and elevation. Our results showed that the relative importance of the site-effect is marginal and accounts for less than 3% of the total information captured (Figure 4). This suggests that the spectral signal from NEON data is comparable across different flights and that flight-specific noise can be minimized using BRDF corrections and vector normalization to limit the impact on the accuracy of generalized algorithms. This is due in part to NEON data being highly standardized and using the same image pre-processing protocol across the entire network (Kampe et al., 2014). NEON remote sensing data is also collected at the peak of vegetation productivity for each site, reducing the confounding effect of different phenological stages for species occurring at multiple sites (Gartner et al., 2016). Expanding large scale surveys outside the NEON network would require integrating information from less standardized sources, raising new challenges related to fusion of sensors that are not cross-calibrated and images collected in different seasons (Brook & Ben-Dor, 2015, Zou et al., 2018). Further investigation is therefore fundamental to evaluating whether our findings apply to applications that involve integrating multiple sensors, missions, or resolutions.

Clustering adjacent bands in the electromagnetic spectrum using KLD facilitated evaluating tree attributes, such as leaf chemistry, that may allow spectral separation of different species. The phylogenetic conservation of these attributes may help explain why a large part of the confusion in species classification was for congeneric species (Cavender-Bares et al., 2016). Our results indicating important spectral regions support patterns shown in previous work, including (a) reflectance in the red edge (Curran et la., 1995), (b) 450-475 nm (Kira et al., 2015), and (c) the SWIR around 1200 nm (Li et al., 2021), 1600 nm and 2000 nm (Kokaly et al., 2015). The importance of the 450-475 nm region may be linked to carotenoids and chlorophyll content, with chlorophyll content generally lower in needleleaf species (Croft et al., 2020) and carotenoids varying across different environments (Valiente et. al, 2015). Reflectance in red-edge can be related to leaf age, chlorophyll, and pigment concentration (Gitelson et al., 1996) that vary widely among species (Cavender-Bares et al., 2016). Reflectance in the 1200 nm was previously linked to equivalent water thickness (Li et al., 2021), a key functional trait for classifying species in temperate biomes (Shi et al., 2018), or distinguishing early to late succession species (Feret et al., 2019, Wright et al., 2004). Reflectance in SWIR at 1600 and 2000 nm can be linked to leaf phenolics (Kokaly et al., 2015), tannins and secondary metabolites (Couture et al., 2016), proxies of leaf toughness and structure across species. The link between water thickness, toughness and structure may also explain why the regions in 1200 nm and 1600 nm are the two most important in distinguishing broadleaf from needleleaf species.

The dimensionality reduction algorithm used in this study identified groups of adjacent bands in relatively discrete spectral regions that overlap with spectral regions used in multispectral satellites, supporting the idea that multispectral satellite sensors can access a large amount of spectral information for species classification (Laurin et al., 2016). Hyperspectral satellite data is still limited to few prototype datasets with relatively low spatial resolution (Loizzo et al., 2018, Diaz et al., 2018, Bogan et al., 2019), compared to multispectral satellites with sub-meter resolution (e.g. WorldView3). Our results show that most of the information required for species classification across NEON sites overlap with WorldView3 satellite multispectral bands supporting that species identification at the tree and plot level with satellite data is feasible (Immitzer et al., 2012, Hartling et al., 2019, Ferreira et al., 2019).

One of the advantages to broad scale general models is that they allow assessment of how different ecological and environmental conditions influence the accuracy of the species classification. Understanding variation in model performance across space, forest types, and taxa is fundamental to better understanding where and when these models can be applied and improvement of large-scale surveys from remote sensing. In our analysis, eastern US forests showed lower accuracy compared to western ecosystems. We believe this is at least partly because eastern ecosystems are characterized by a higher species diversity of canopy trees as well as crown geometry that makes aligning stems to crowns more difficult compared to western conifer stands (Figure 2, 3, S.6). Higher species diversity in eastern forests (mean species per site ∼15) compared to western forests (mean species diversity per site ∼4), inherently makes classification tasks more challenging due to larger numbers of classes typically resulting in lower accuracy predictions (Takahashi et al., 2020). Continuous closed canopies also increase the likelihood pixels selected in a window centered on the stem will be from neighboring tree crowns. This is due to the difficulty of obtaining accurate GPS points of stems in closed canopy (Rodriguez-Perez et al., 2007), as well as the increased likelihood of sunlit portions of the crown being displaced from the stem location in continuous broadleaf forests (Strigul et al., 2008). This is a common problem, since field surveys often provide only the geographic coordinates of tree stems and lack information about crown position or size, making it very challenging to correctly align crown borders with species labels. For example, pixel mislabeling may be one of the reasons why our classifier was weaker at sites in the Great Plains region (e.g., the NEON sites of Lyndon B. Johnson National Grasslands, CLBJ and University of Kansas Field Station, UKFS), where patches of grasslands alternate with dense forests characterized by multiple oak species forming a complex mosaic of crowns that may not be located directly above their stem locations. In contrast, conifers in western US forests tend to be dominated by species characterized by apical dominance (e.g., aspens and firs) with crowns centered directly above the main stem, reducing pixel mislabeling and improving classification. Finally, savannas, such as the San Joaquin Experimental Range (SJER), characterized by isolated trees of few species (mostly broadleaved), may be less likely to suffer from confounding effects like crown displacement and stem-crown misalignment, making them less prone to spectral mixing or potential pixel-mislabeling (Heinzel & Koch, 2012). The most challenging ecosystem type in our analysis, wetlands, combines all of these challenges. Species like *Carpinus caroliniana* and *Betula papyrifera*, found often in plots from wetland ecosystems, were among the species with the worst classification accuracy, partly because they are generally smaller trees that can occur in the understory, grow in closed canopies in the overstory (e.g., an average dbh ∼16.5 cm and average height of ∼10 m), and often include limited training samples because they are mostly found in riparian ecosystems which make up a small fraction of the landscapes from the NEON sites included in this study (less than 50 individuals per species). Because of these challenges, the accuracy of the species predictions needs to be assessed depending on the site and ecosystem types within sites to ensure it is sufficient for the intended ecological application.

Because species predictions from remote sensing are imperfect, it is important that classification models produce robust estimates of uncertainty to allow this uncertainty to be propagated through ecological analyses and considered during decision making. This is particularly important when generating large numbers of predictions at large scales, because this will result in including species located in undersampled areas and challenging ecosystems as well as species that are difficult to classify due to rarity or similarity to other closely related species. Our results confirmed that stacking scores from different classifiers using a logistic regression produces accurate estimation of classification uncertainty (Figure 6).

While our generalized approach resulted in significant improvements over site-level models, it is important to recognize that the accuracy of this approach is still insufficient for ecological analyses contingent on rare or untrained species. For example, biodiversity patterns are often driven by rare species (Leitao et al., 2016, Mouilllot et al., 2013), which are the most challenging taxa for species classification, especially species so rare that they cannot be included in the model due to data limitations (n < 5 individual trees in this study). Extrapolating outside of NEON sites, a goal for general models, would also result in the presence of additional species missing from the field dataset, restricting the range of ecological analyses to species sampled within the footprint of NEON sites. Some of these limitations may be mitigated by classifying trees into higher level taxonomic levels. In this study we observed that misclassified trees were generally limited to species in the same genus and species co-occurring in the same plot (Figure S.9, Figure S.10, Figure S.11, Supplement 3). Oaks, pines, and poplars in particular accounted for ∼87% of the total within genus confusion. One possible driver of confusion among oak species is their similar physiological and spectral characteristics (Figure S.12). Some co-occurring oaks species like *Quercus alba* and *Quercus stellata* can also cross-breed and therefore be physiologically very similar (Hardin, 1975), making them particularly hard to distinguish from imagery. For these reasons, most of the cases leading to misclassification resulted from within-genus confusion. This implies that uncertainty can be significantly reduced by aggregating predictions to the genus level, offering a more robust solution for large scale ecological applications that can be successfully addressed by accurately classifying trees at the level of genus, families, or plant functional type. For example, earth system models use plant functional types as the taxonomic unit for quantifying carbon dynamics at continental to global scale (Lawrence et al., 2019), large scale fire risk assessment and management can be achieved by using genus level surveys of the most dominant taxa (Ma et al., 2021), and patterns of forest biomass largely depend on which taxa dominate the ecosystem (Cheng et al., 2018). An increase in taxonomic level also reduces issues with applying general models beyond the training data (e.g., outside of NEON sites) because it is much more likely that all genera or families have been sampled in the training data.

Building generalized algorithms provides an approach to overcome the significant field data limitations present in most remote sensing tasks in ecology, by allowing pooling data from ever growing sources of spatially explicit field surveys and high-resolution remote sensing imagery. Our results showed that by integrating field surveys from dozens of NEON sites, it is possible to produce a general model that provides improvements over single-site models for species classification, with good estimates of uncertainty, and the ability to increase accuracy further by aggregating predictions at the genus level. This general approach also unlocks the potential for making predictions outside of NEON sites. The ultimate goal is to develop general models that can be used anywhere in the region of interest (in our case the United States). Using only NEON data, we successfully built a single integrated classifier that includes 20% of all tree species found in forest ecosystems across the US (n=77 out of 396 surveyed by the United States Forest Inventory and Analysis project; appendix F; Woudenberg, et al., 2010).

Beyond NEON, more and more openly available field, multispectral and hyperspectral datasets are being released from aerial (airborne and UAV) and satellite missions worldwide (Cook et al., 2013, Vangi et al., 2021, Claverie et al., 2018). Our results show that instead of training hundreds of separate models for local applications, there is the potential for integrating field and remote sensing collections from multiple locations and sources to build general models with improved accuracy for a broader range of landscapes and geographic locations. Leveraging the broad geographic distribution of NEON sites and the overlapping information held by multispectral and hyperspectral imaging, our results also suggest the potential for linking different data sources to unlock the ability of scaling species classification of individual trees beyond NEON. For example, future work could focus on developing approaches for bridging the information held in hyperspectral data (sparsely acquired, high radiometric resolution) to the ever-growing pool of high-resolution multispectral and RGB + NIR data (e.g. National Agriculture Imagery Program data) available for a broad geographic continuum across the US. Further integration with more field and remote sensing datasets could potentially provide remote sensing-based surveys of hundreds of millions of trees, making it possible to investigate the properties of ecosystems from local to continental scales.

## 5. Conclusions

Remote sensing is facing a revolution in the quality of data and accuracy of methods, making it a good candidate for developing applications to survey species and forest properties at large spatial extents. Leveraging data collected from NEON across the US, we demonstrated that building continental scale algorithms for generalized species classification offers several advantages over the more traditional site level applications. Despite being very high for dominant taxa, accuracy in predictions for less represented species can be taunted by limitations in field-to-image misalignment, the number of species and individuals from rarely sampled taxa, making surveys from remote sensing unsuited to date for analyzing patterns in species alpha diversity at scale. Yet, building generalized algorithms is a fundamental cornerstone to overcome these limitations, because it allows for pooling from ever growing sources of geo-explicit field surveys and high-resolution remote sensing imagery. Our results showed that by integrating field surveys with NEON airborne data, it is possible already to generate highly accurate predictions at the genus level and overall good estimates of uncertainty for individual trees. This allows for generating surveys of hundreds of millions of individual crowns across the continent, unlocking the potential for investigating large scale ecological applications focusing on the sun-exposed part of the canopy, dominant species, genuses or functional types.

## Acknowledgements

This work was supported by the Gordon and Betty Moore Foundation’s Data-Driven Discovery Initiative through grant GBMF4563 to E. P. White and by the National Science Foundation through grant 1926542 to E. P. White, S. A. Bohlman, A. Zare, D. Z. Wang, and A. Singh; by the NSF Dimension of Biodiversity program grant (DEB-1442280) and USDA/NIFA McIntire-Stennis program (FLA-FOR-005470) to S. A. Bohlman; by the University of Florida Biodiversity Institute (UFBI) and Informatics Institute (UFII) Graduate Fellowship to Sergio Marconi. There was no additional external funding received for this study.

All confusion matrixes can be found in the supplementary material. All data can be found in the following Zenodo archive: https://doi.org/10.5281/zenodo.5796143. All code for data preprocessing, model training and testing and analyses can be found in the following GitHub repo: https://github.com/MarconiS/Continental-scale-Hyperspectral-tree-species-classification-in-the-National-Ecological-Observatory-N

Supplement 1: parameterization of classifiers and meta ensemble

KNN classifier was trained using 20 neighboring points, with distance weighted by the inverse of their Manhattan distance. The Random Forest classifier was trained using 300 trees with up to 7 features (square root of the total predictors) considered for better split, validated using cross-entropy loss function on out-of-bag samples. The gradient boosting classifier was trained using 1000 maximum iterations, a learning rate of 0.01, max depth of 25 and 0.5 L2 regularization. Loss was calculated using categorical cross-entropy on out-of-bag samples. Multi-layer Perceptron classifier was trained for 1200 max iterations, using relu activation, 1 hidden layer, weight optimization through adam booster with exponential decay rate of 0.9. The Bagging Classifier was trained using 10 support vector machine classifiers as base estimators. We used loose regularization (C = 1000), RBF kernel, and 5-fold cross validation to calibrate probability estimates. The meta ensemble was trained using probability vectors produced by each weak classifier. We used a regularized logistic regression (elasticnet), with 0.5 L1 to L2 penalty ratio. We used a saga solver to optimize the loss function.

Supplement 2: Species level accuracy and scientific names

Species names for all species used for this manuscript along with their precision, recall and accuracy can be found in the supplementary file titled “overview_precision_recall_names.csv”. Recall is defined as the amount of true positives divided by the sum of true positives plus and false negatives; it represents the fraction of relevant instances predicted by the model. F1 represents the model accuracy for each species.

Supplement 3: Confusion matrix

Confusion matrices were produced using the Caret R package (Kuhn, 2008). For species with both precision and recall equaling 0, F1 score was calculated as 0. Tabular version of the confusion matrices for predictions on the test (total n = 1487) set for (1) all trees in the test set, (2) trees in the test set for each ecodomain, (3) trees in the test set for each site from the generalized approach, (4) trees in the test set for each site from site-specific approach, (5) for predictions at the genus level can be found in the supplementary file “confusion_matrices.zip” and are organized in separate folders. For each confusion matrix, rows represent observations, columns represent predictions. In cases where columns are entirely filled with zeros, we removed all species that were not found in either the training or held-out test datasets at each individual site. For site level confusion matrices, we only included species for which at least one tree was either observed or predicted. Therefore, species with no observations in the test set will be assigned to empty columns; species never predicted in the test set will be assigned to empty rows. This applies to the overall confusion matrix too, where tested trees were mis-predicted as Gleditsia triacanthos and Quercus michauxii despite these two species not being included in the held-out test dataset (empty rows).

## Supplementary Figures

**Figure S1.**
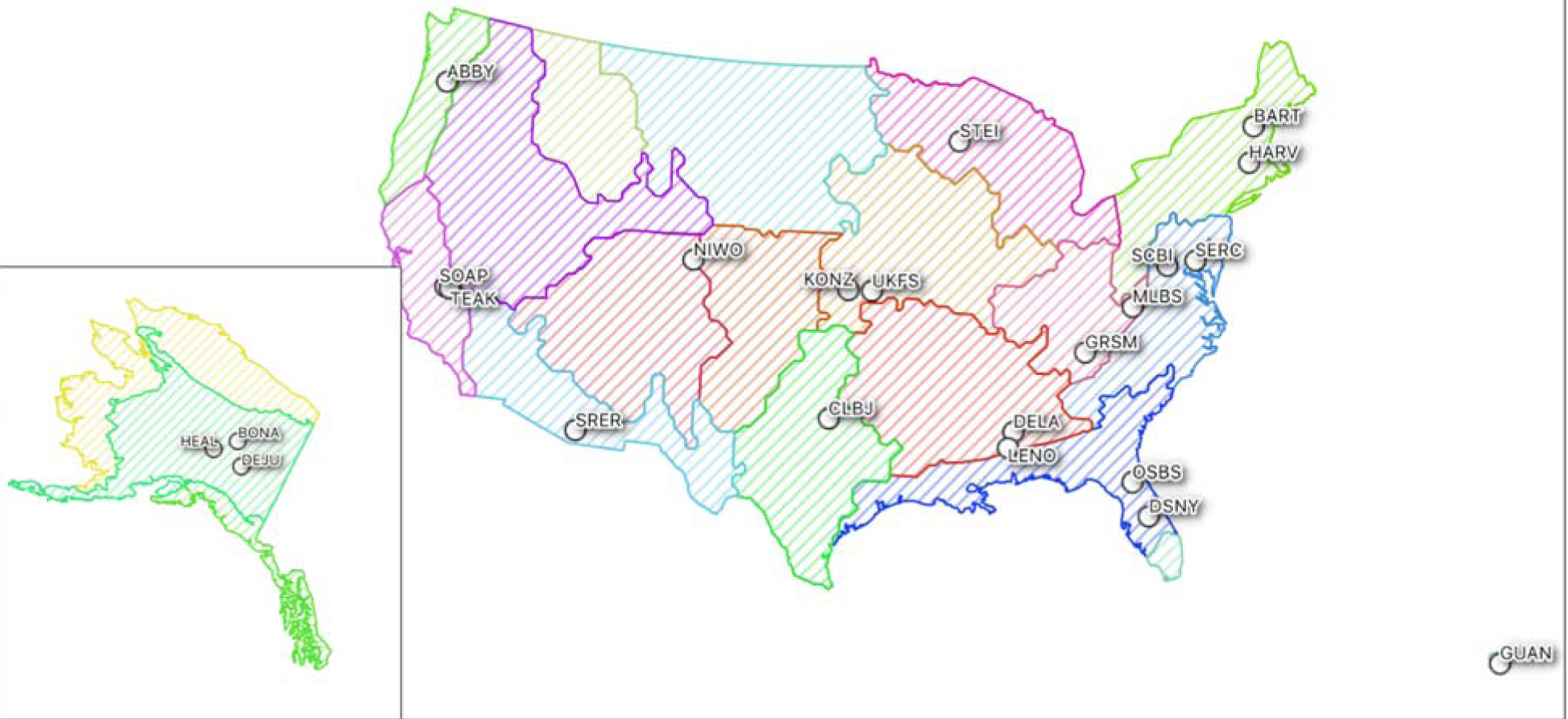
Geographic distribution of NEON sites included for this study. Colored regions represent ecological regions defined by NEON (https://www.neonscience.org/field-sites/about-field-sites). A description of each site and their ecological domain can be found in the Supplementary Table 1.

**Figure S. 2:**
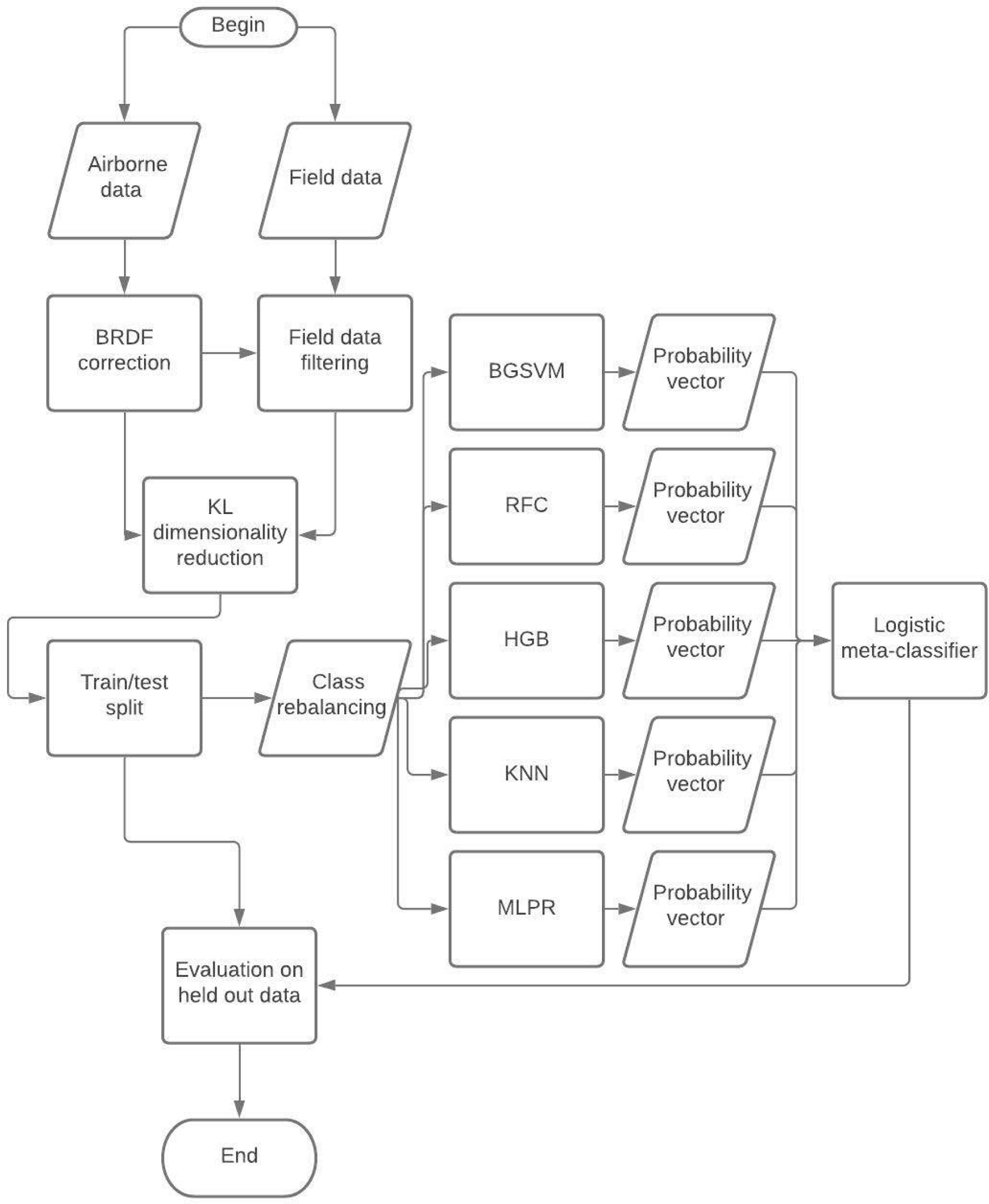
Flowchart of the species classification pipeline developed for this study

**Figure S.3.**
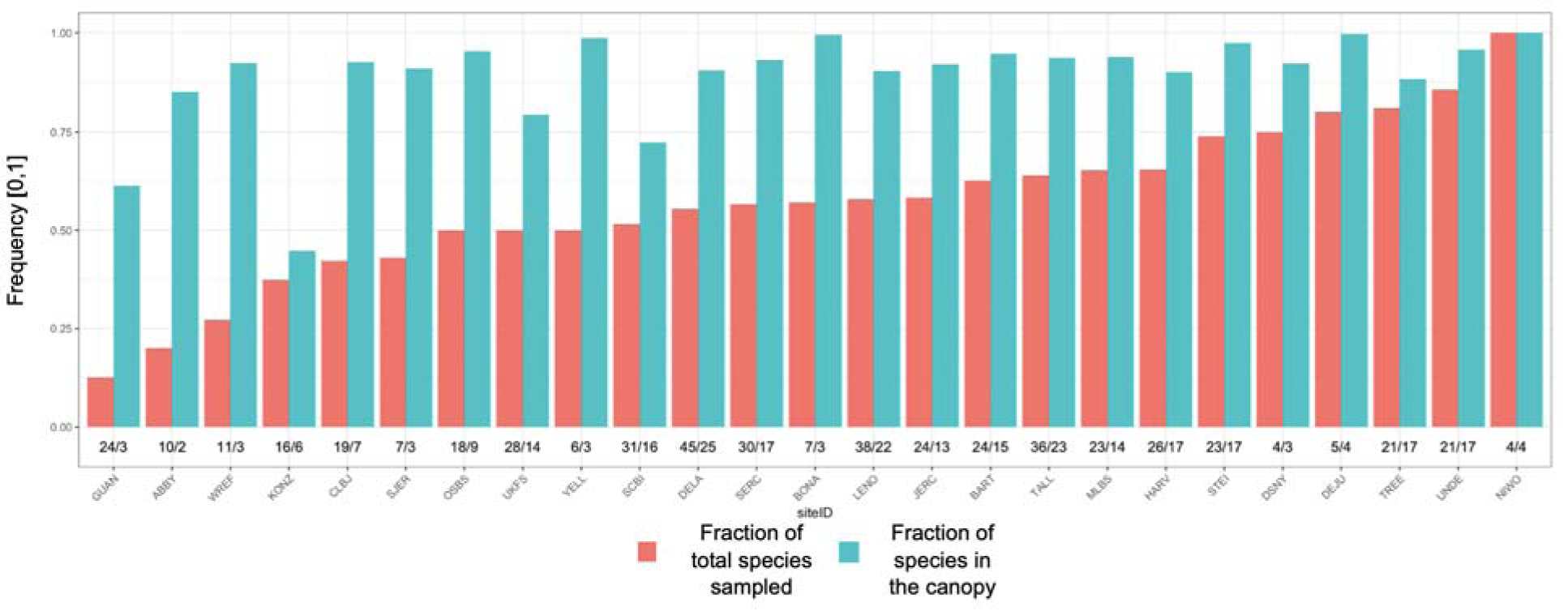
For each site, the fraction of species included in the test/train dataset compared to the total amount of tree species in the raw NEON vegetation structure dataset (red); the fraction of trees that the species from the test/train dataset comprise out of all canopy trees (blue) in the NEON vegetation structure dataset. The numbers separated by “/” above each site name represent the total number of species in the original dataset and in the filtered data respectively, specific for each site. Trees in the canopy (blue bars) were determined by canopy position data in the vegetation structure data where trees in the canopy were designated as “Full sun”, “Mostly shaded”, “Partially shaded”, “Open growth”, or non-classified (“NA”).

**Figure S.4.**
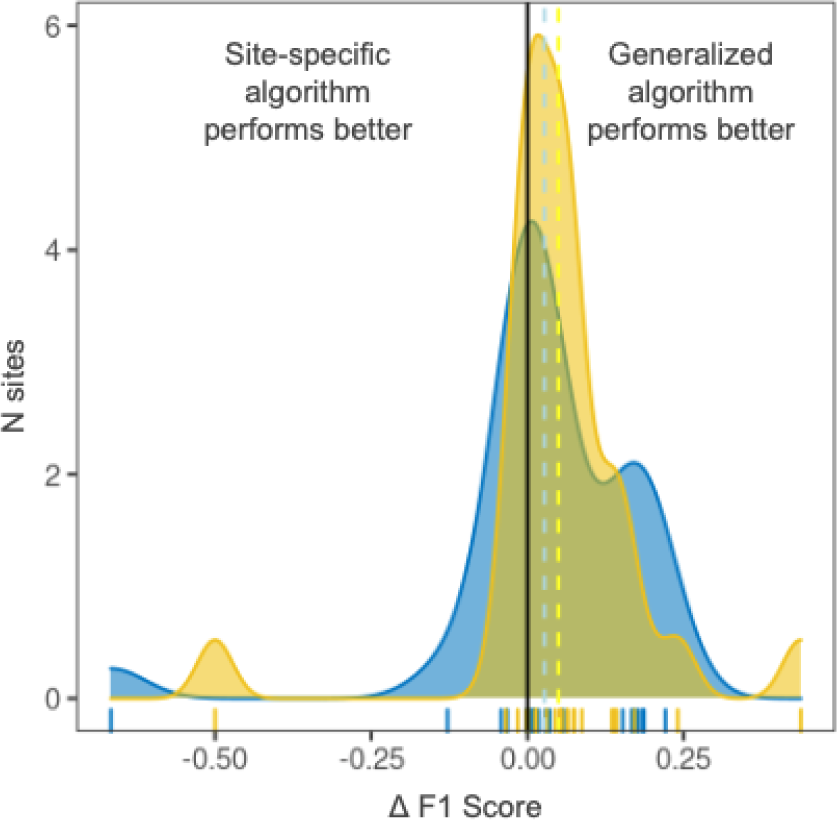
Density functions of the difference in ΔF1 scores between the generalized and each single-site algorithm for species-level F1 (yellow) and individual-level F1 (blue). Positive ΔF1 values (17 out of 27 sites) represent sites where the generalized algorithm outperformed its site specific counterpart. Dashed vertical lines represent the average ΔF1 across sites (species-level F1 = 0.09, individual-level F1 = 0.05).

**Supplement S.5.**
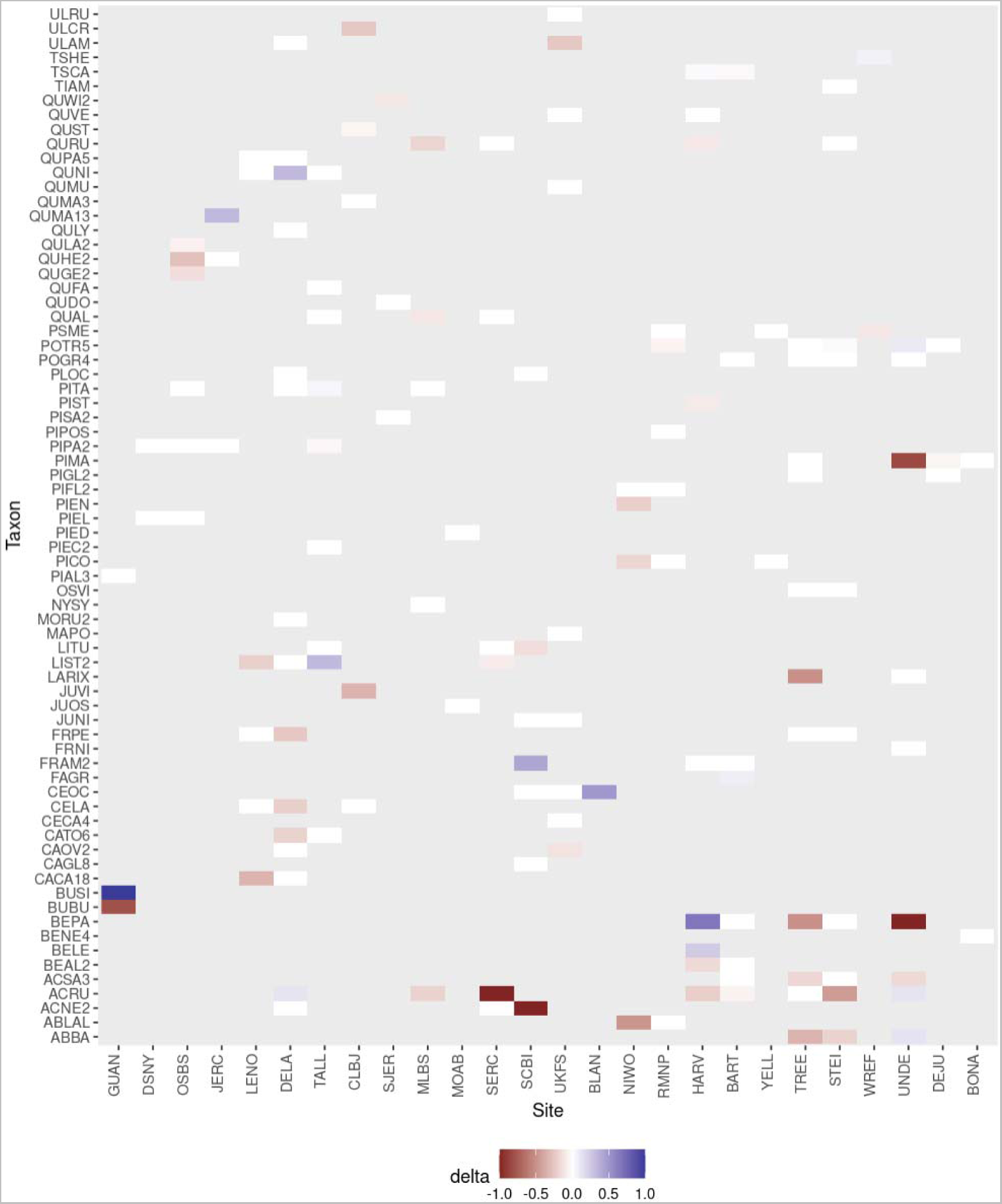
Difference in accuracy between the general and site-specific approaches for each species-site combination. Negative values (red) represent taxa whose accuracy is higher in the general approach. Blue values represent taxa whose accuracy is higher in the site-specific approach. White values where accuracy was similar for the general and site-specific approaches. Grey are species that do not occur at the site. Sites are sorted by geographic similarity. Species names for each taxon acronym can be found in Supplement 2. Site names can be found in table S1.

**Figure S.6.**
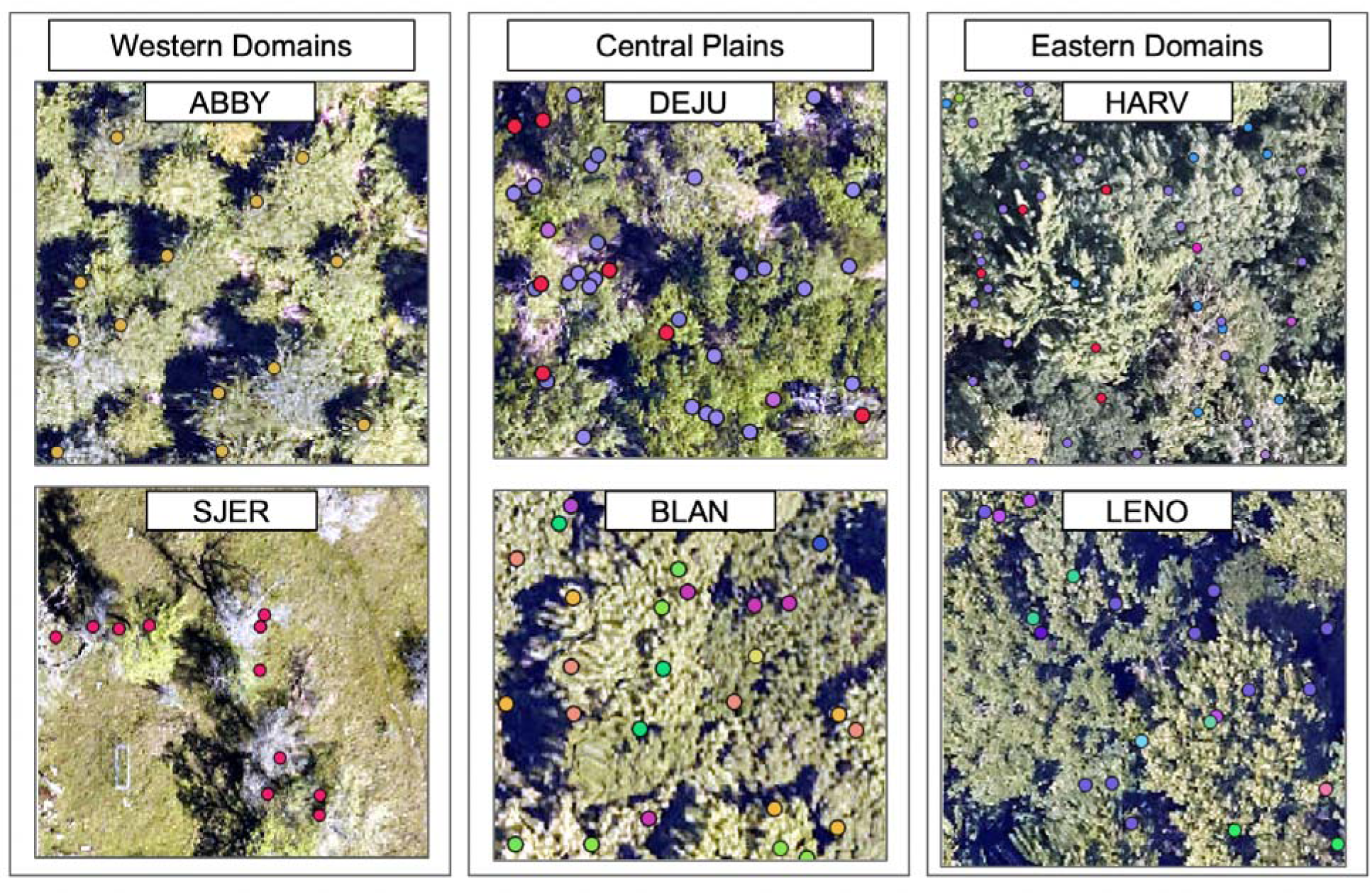
Example of 400 m^2^ plots for 6 sites from western (left panel), central (center panel) and eastern US (right panel). Dots represent field stem data collected from NEON vegetation structure. Different dot colors represent different species. Only stems that have been filtered to include only stems that are likely to be in the canopy. From top left to bottom right sites acronyms are Abby Road (ABBY), Delta Junction (DEJU), Harvard Forest (HARV), San Joaquin Experimental Range (SJER), Blandy Experimental Farm (BLAN), Lenoir Landing (LENO),

**Figure S.7.**
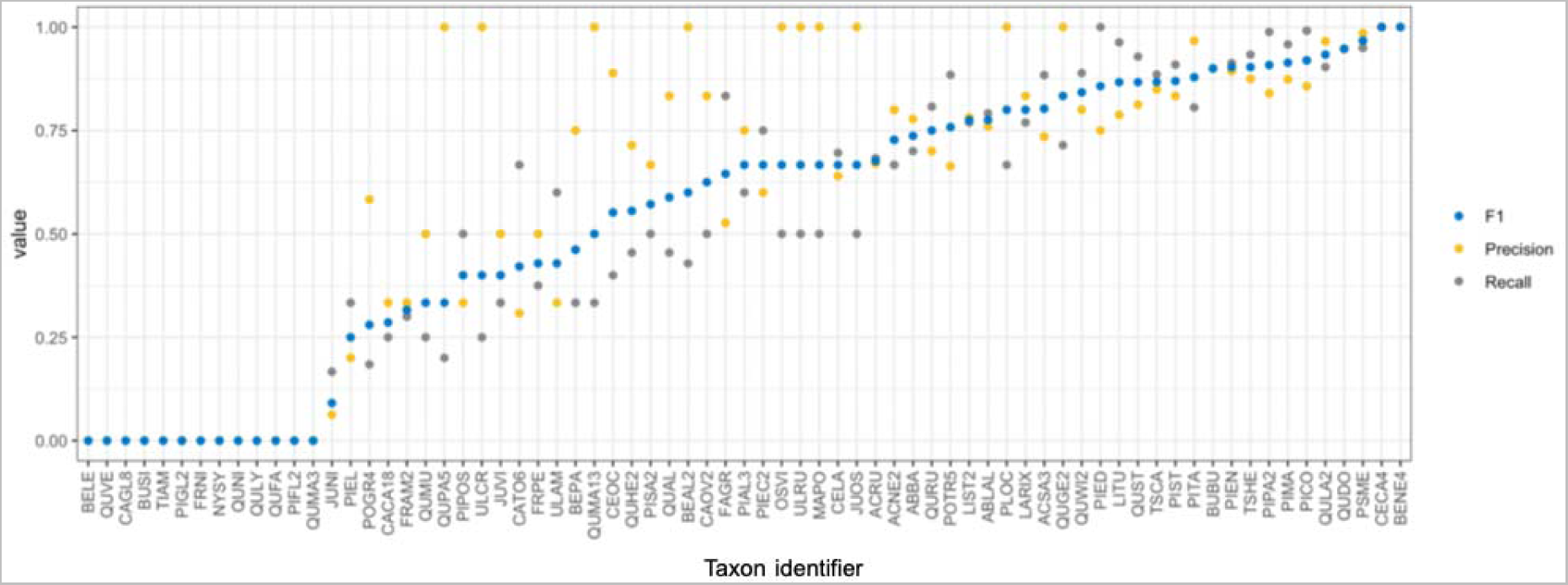
Precision (blue), Recall (yellow) and F1 (gray) for each individual species included in the dataset. Precision is defined by the ratio between the number of true positives and the number of true positives plus the number of false positives; it represents the ability of a classification model to identify only the relevant data points. Recall is defined as the amount of true positives divided by the sum of true positives plus and false negatives; it represents the fraction of relevant instances predicted by the model. F1 represents the model accuracy for each species. These results, along with the list of species scientific names assigned to each code can be found in Supplement S2. Confusion matrices can be found in separate supplementary file as described by Supplement 3.

**Figure S.8.**
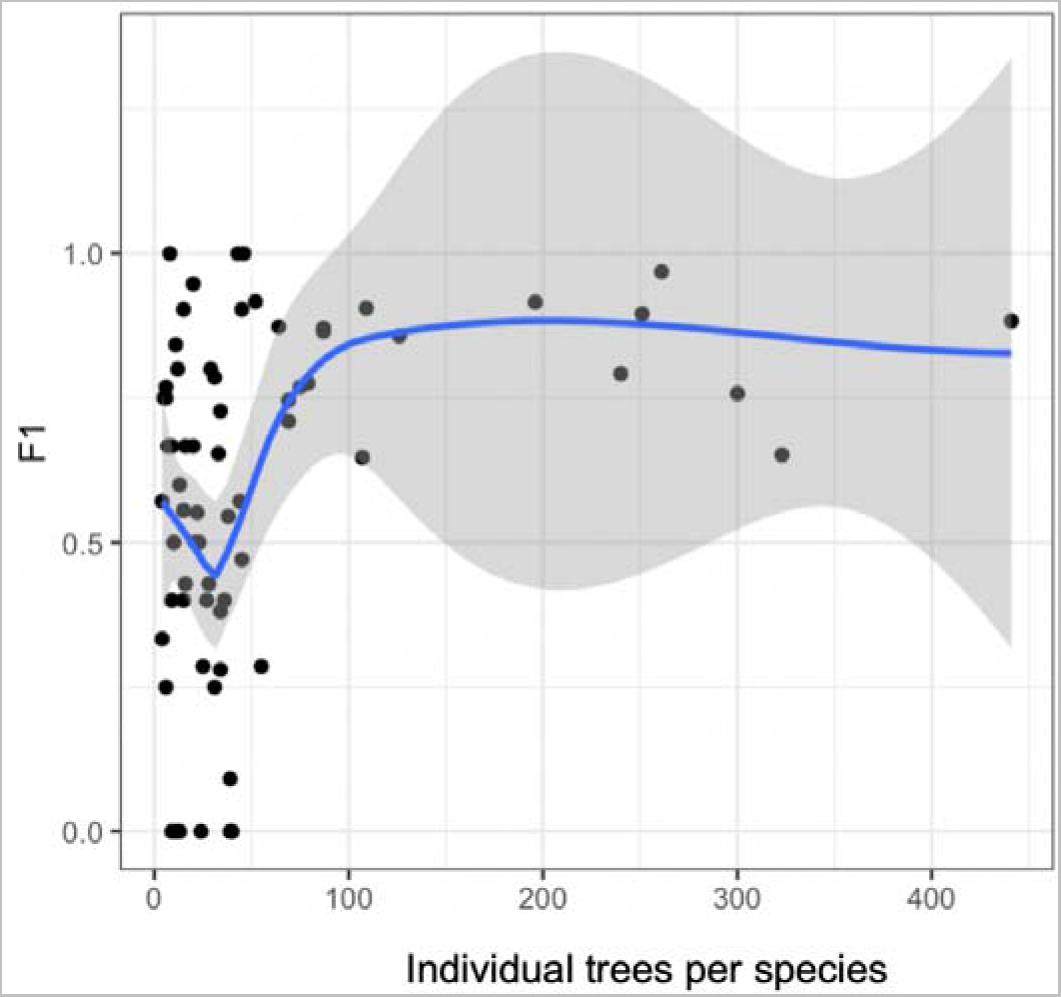
The relationship between individual species F1 scores and number of individual trees available for training for that species. The blue line shows a fitted relationship using local polynomial regression fitting (loess) and the grey region shows the 95% confidence interval around that relationship.

**Figure S.9.**
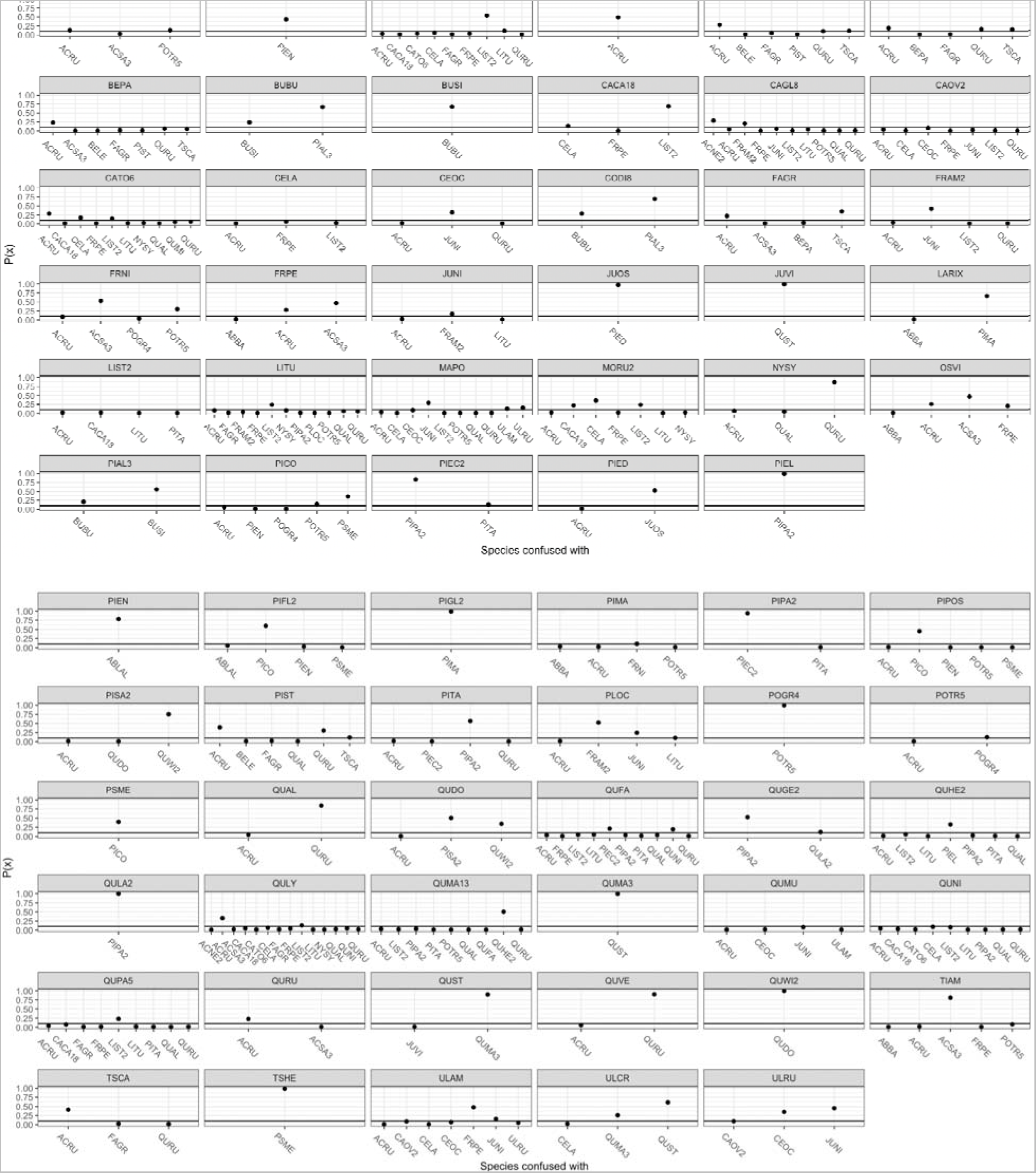
Confidence score P(x) for those taxa each species was most confused with. Taxa with a P(x) lower than 0.02 were not included. Species names for each taxon acronym can be found in supplement 2.

**Figure S.10.**
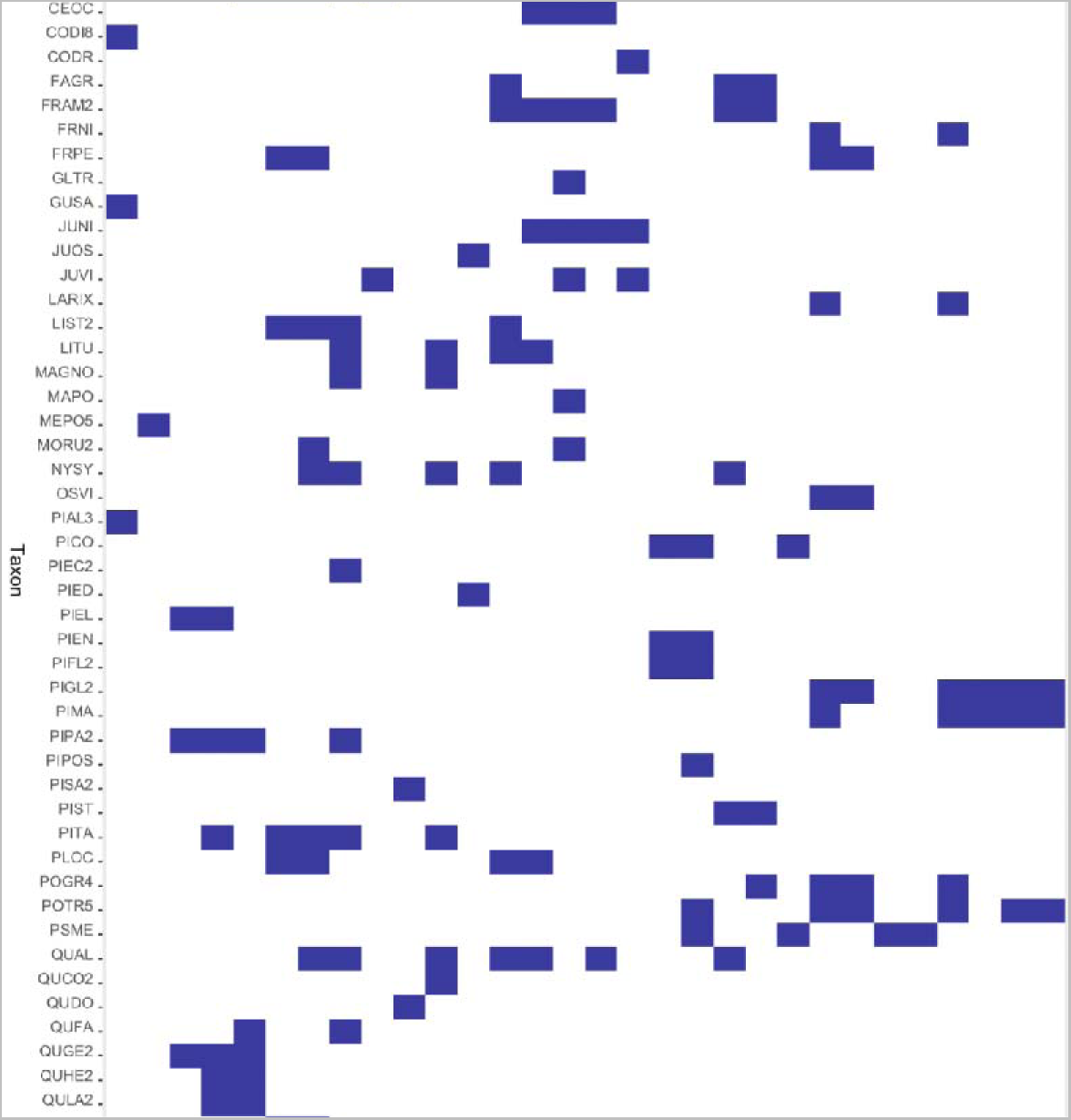
Distribution of species across sites. Species names for each taxon acronym can be found in supplement 2. Site names can be found in table S1.

**Figure S.11.**
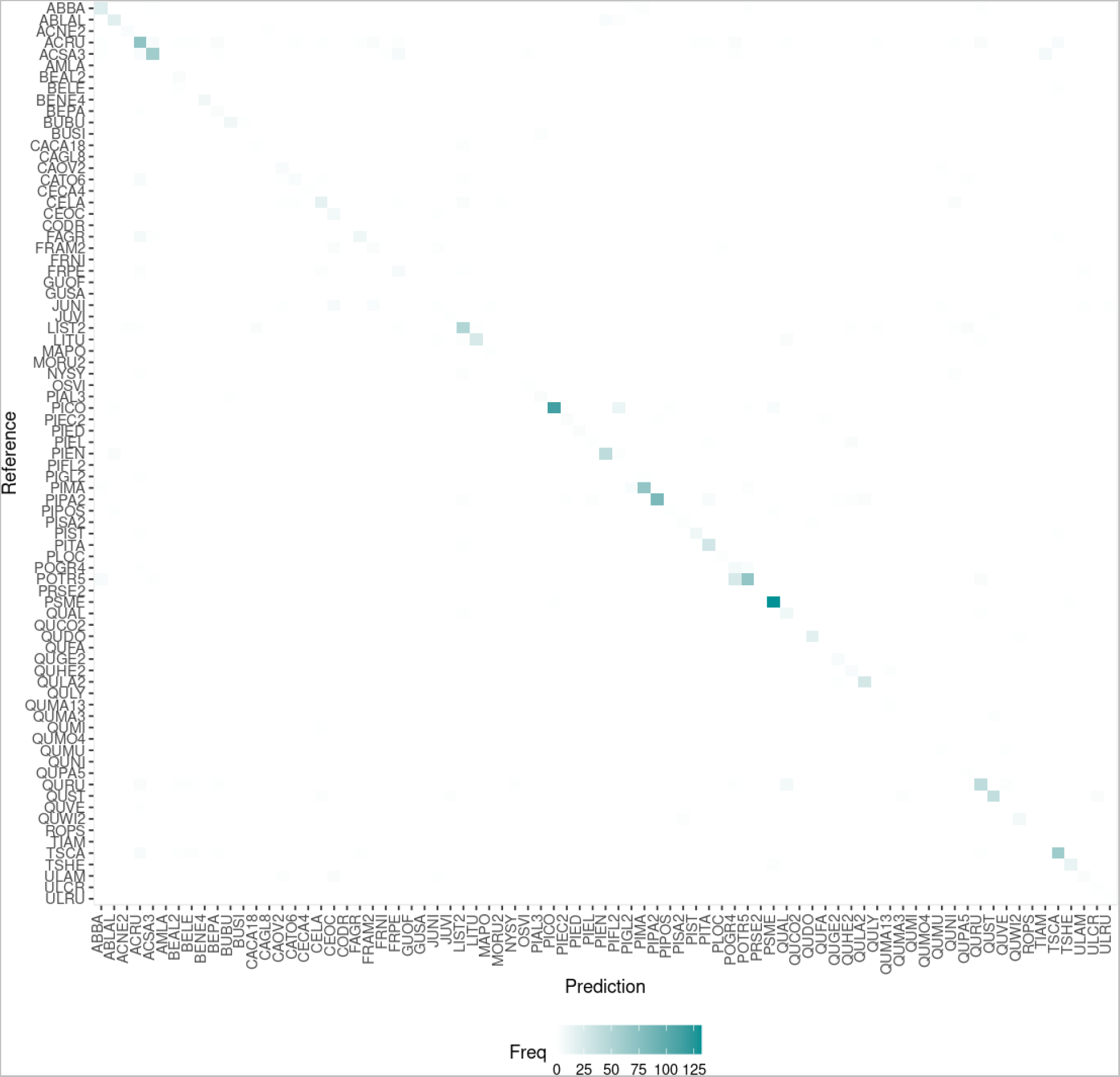
Overall confusion matrix for all data in the test set. Tabular versions, including the confusion for each site, ecodomain and the confusion matrix for predictions at the genus level can be found in the supplementary files. Species names for each taxon acronym can be found in supplement 2.

**Figure S.12.**
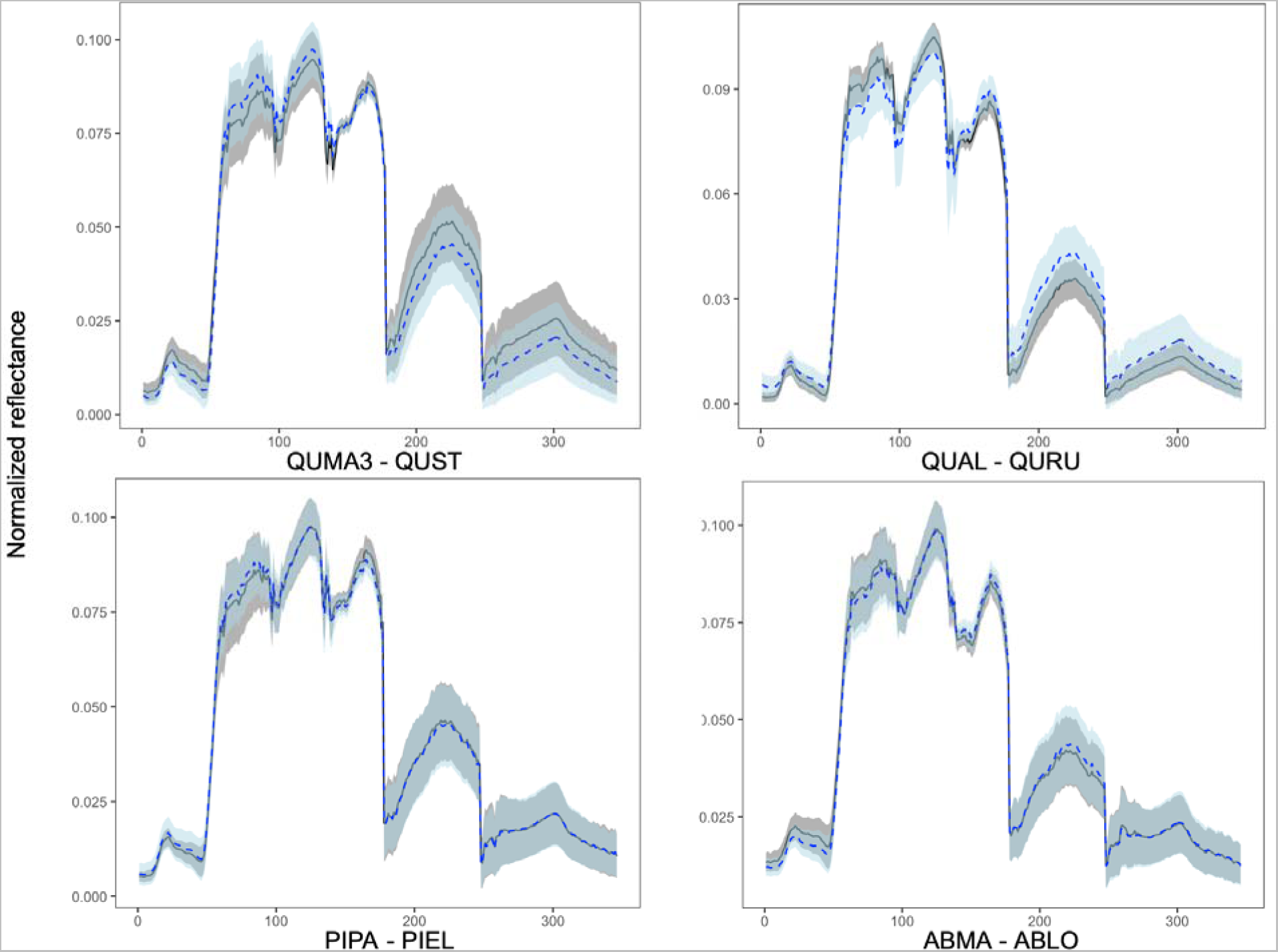
Example of spectral signature overlap between often confused congeneric species. Lines represent the average spectra, shaded areas represent the standard deviation for all pixels extracted for that particular species. The first species in legend is in blue, the second in grey. The X-axis is the band number from brdf corrected hyperspectral image. Couples of species are: (a) *Quercus marilandica* and *Quercus stellata*, (b) *Quercus alba* and *Quercus rubra*, (c) *Pinus palustris* and *Pinus elliottii*, (d) *Abies magnifica* and *Abies lowiana*.

**Table S.1.**
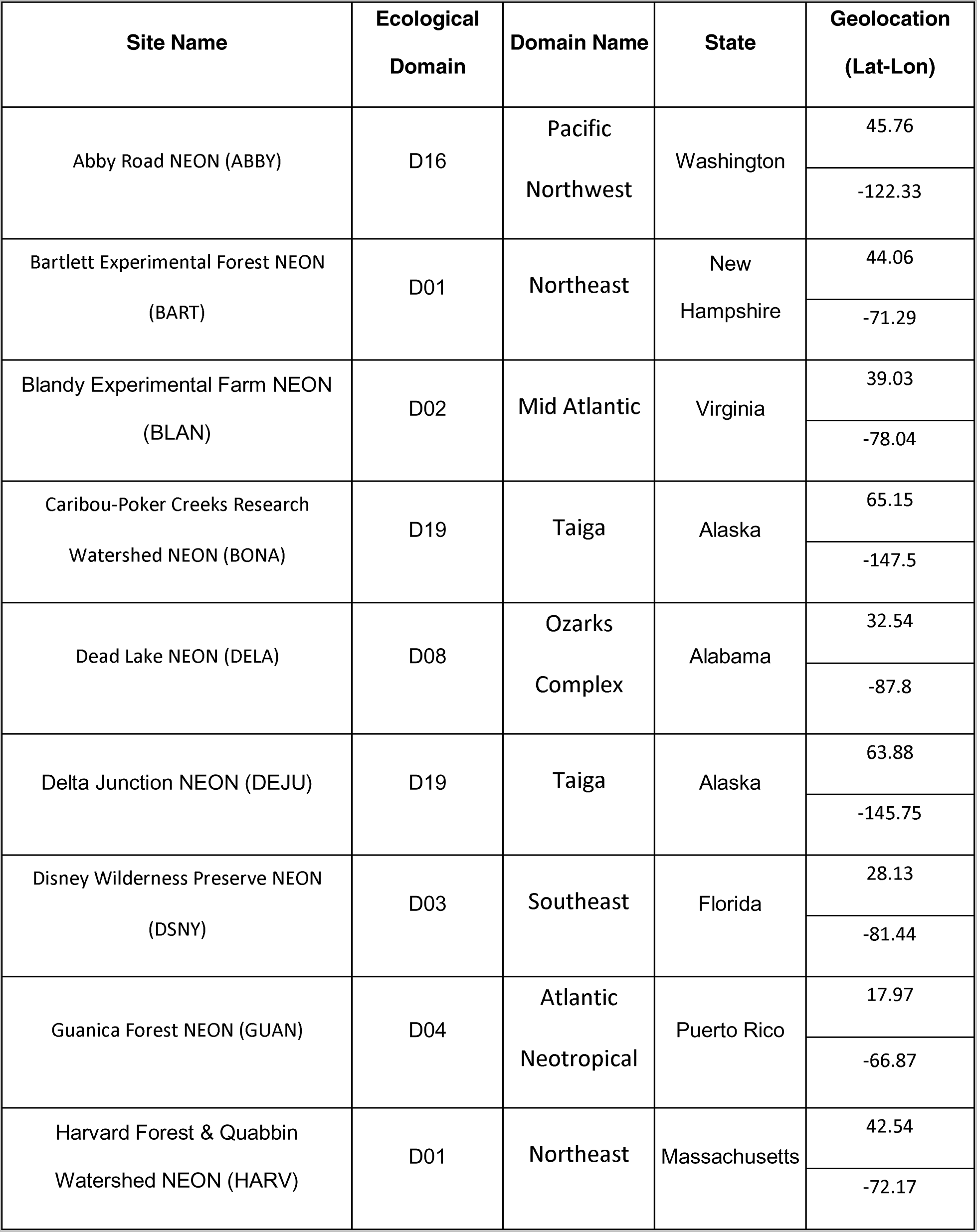
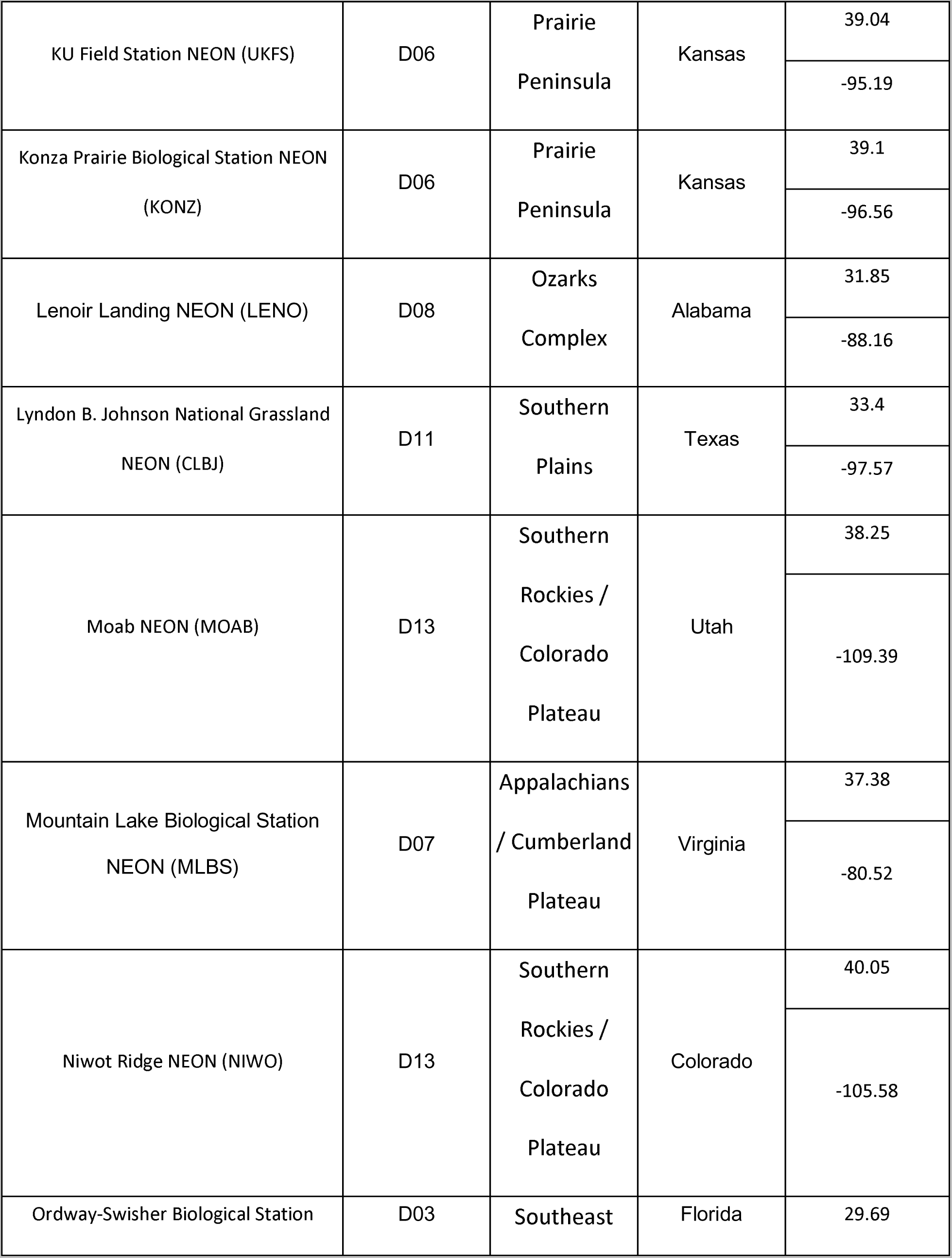
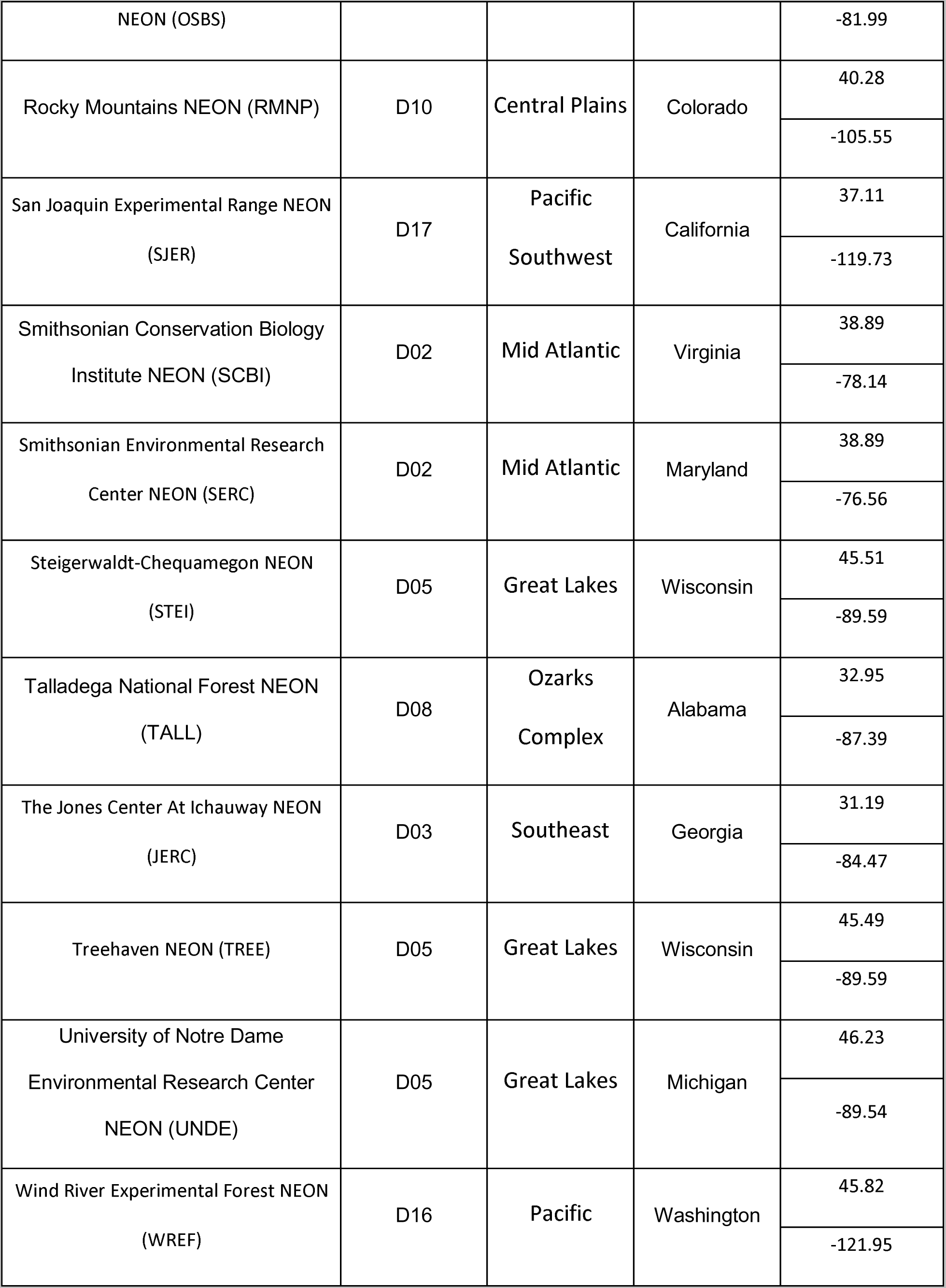
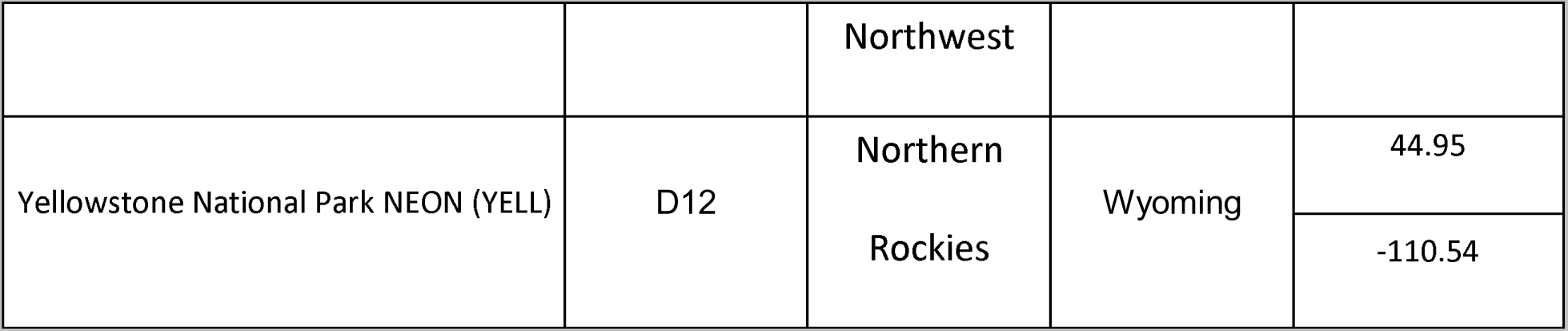
Description of NEON sites and ecological domains used in this study.

